# Co-cultivating rice plants with *Azolla filiculoides* modifies root architecture and timing of developmental stages

**DOI:** 10.1101/2024.10.02.615985

**Authors:** Sara Cannavò, Chiara Paleni, Alma Costarelli, Maria Cristina Valeri, Martina Cerri, Antonietta Saccomanno, Veronica Gregis, Graziella Chini Zittelli, Petre I. Dobrev, Lara Reale, Martin M. Kater, Francesco Paolocci

## Abstract

Strategies for increasing the yield of rice, the staple food for more than half of the global population, are needed to keep pace with the expected worldwide population increase, and sustainably forefront the challenges posed by climate change. In Southern-East Asian countries, rice farming benefits from the use of *Azolla* spp. for nitrogen supply. In virtue of the symbiosis with the nitrogen-fixing cyanobacterium *Trichormus azollae, Azolla* spp. are ferns that release nitrogen into the environment upon decomposition of their biomass. However, if and to what extent actively growing *Azolla* plants impact on the development of co-cultivated rice plantlets remains to be understood. Here, we show that actively growing *Azolla filiculoides* plants alter the architecture of the roots and accelerates the differentiation and proliferation of leaves and tillers in co-cultivated rice plants. These changes result from an intimate cross-talk between rice and *A. filiculoides,* in which hormones and other metabolites released by the fern in the growth medium trigger an alteration in the rice root transcriptome and the hormonal profiles of both roots and leaves. Overall, the present data let us argue that co-cultivation with *A. filiculoides* might prime rice plants to better deal with both abiotic and biotic stress.

**Highlight:** *Azolla filiculoides* alters the root transcriptome and hormonal balance in both roots and leaves of co-cultivated rice plantlets, thereby interfering with the progression of their developmental programs

## Introduction

Rice (*Oryza sativa*) is a staple food crop for more than half of the global population, which is expected to reach almost 10 billion in 2050 (UN 2022). As global demand for major crops will roughly double by 2050, agricultural production may need to be increased by 70%–110% (FAO, 2009a; Tilman *et al*., 2011). Unfortunately, the increase in the size of the human population is paralleled neither by the yield increase of rice nor by any other agricultural products (Godfray *et al*., 2010; Alexandratos and Bruinsma, 2012; FAO, 2009a, b, 2022a, 2022b). The stagnation of crop yields has several underlying reasons, including loss of genetic diversity in today’s crop species (Day, 1973; Gaillard *et al*., 2018; Mueller *et al*., 2012). Moreover, the increasing temperature, rising sea levels, and alterations in rainfall patterns and distribution caused by global climate change could lead to substantial modifications in land and water resources for rice production as well as in the productivity of rice crops grown in different parts of the world (Nguyen, 2002). Therefore, the ongoing climate change poses a further obstacle to assuring food security and achieving the most significant Sustainable Development Goals (SDGs) set by the UN (https://sdgs.un.org/goals).

Rice yield decline has also been related to decreased physiological nitrogen (N) use efficiency (NUE). This occurs in intensive rice cultivation systems under tropical conditions and when rice is grown under temperate conditions, where N is supplied as urea or green manure (Lhada *et al*., 2000; Casanova *et al*., 2002). Yet, in Southern-East Asian countries, the aquatic ferns belonging to the *Azolla* genus have been traditionally used in rice farming as N supplier crops either as a mono-crop, when they are incorporated into the soil before rice transplanting, or as intercrop, when grown as a dual crop along with rice (Fogg *et al*., 1973; Lumpkin and Plucknett, 1980; Shi and Hall, 1988; Watanabe, 1982; Watanabe and Liu, 1992; Yang *et al*., 2018). Depending on the cropping system, the presence of the fern can increase rice productivity by 30-50% (Marzouk *et al*., 2023), and this increase has been mainly related to increased NUE. *Azolla* spp. are in fact perennial, monoecious floating freshwater ferns that live in a permanent mutualistic symbiosis with the nitrogen-fixing cyanobacterium*Trichormus azollae* and other species-specific endophytic prokaryotic strains. The association *Azolla-T. azollae* is capable of fixing N_2_ at a rate that rivals that of *Rhizobium*-legume symbiosis (Lumpkin and Plucknett, 1980). However, nitrifying bacteria and other plants do not benefit from this low-cost source of N_2_, as the cyanobacteria keep 60% whereas the remaining 40% of the fixed nitrogen is immediately translocated to Azolla (Peters and Meeks, 1989). Only a negligible amount (3-4%) of ammonium (NH_4_^+^) of the total nitrogen fixed by *T. azollae* is released into the water. The remaining 96-97% of the fixed N_2_ is therefore unavailable to other plants until the Azolla biomass is mineralized. This suggests that the Azolla nitrogen-rich biomass is released into the soil only following plant death and decomposition (Mahanty et al., 2017).

Additionally, Azolla considerably modifies the physico-chemical and biological properties of water, controls weed, algae, insects and pest proliferation, thereby reducing resource losses and management costs in rice farming (Herath, 2023). Thanks to these features, Azolla has been recently reconsidered as an eco-friendly and innovative solution to replace or integrate chemical nitrogen fertilizers and pesticides to improve rice yield sustainability even under suboptimal conditions (Yao *et al*., 2018; Khumairoh *et al*., 2018). Although many experiments have demonstrated the increase in the root and shoot growth and, ultimately, weight of grains in rice as a consequence of co-cultivation with Azolla (Bhuvaneshwari and Singh, 2015), the benefit of Azolla on rice varies greatly also according to the climate, the species of Azolla used, and many other factors (Wagner, 1977, Tung and Shen 1985). Studies have documented the competence of both *Azolla* spp. and the endosymbionts hosted in their fronds to synthetise and release metabolites such as siderophores, phytohormones, and volatile organic compounds (VOCs) that might interfere with the developmental patterns of the nearby plants (Vlek *et al*., 2002; Banach *et al*., 2019; Valette *et al*., 2020; Brilli *et al*. 2022). However, the mechanisms underlying the various growth-promoting effects of Azolla on rice are still partially unknown, especially those that may occur during co-cultivation. For instance, the answers to the question of whether Azolla can improve NUE in rice by modifying its root system architecture (RSA) and if it does so not only because it provides N but also via the secretion of growth-promoting substances remain elusive.

The present study focuses on the hypothesis that Azolla is more than a N supplier for rice. In our experimental setup, we cultivated rice plants in the presence of actively growing *Azolla filiculoides* plants to prevent an extra nitrogen supply to rice compared to the control condition (rice plants alone) and gain insights into the morphogenetic effects that *A. filiculoides* has on young rice plants. To reach this goal, the morphological monitoring of rice root apparatus and aerial organs’ development was coupled with rice root transcriptomics and hormonal profiles of rice roots, leaves, and growth media. Our results demonstrate that *A*. *filiculoides* interferes with the patterns and timing of rice root and aerial organs’ development by modulating rice root transcriptional profiles and the hormonal balance in both rice roots and leaves.

## Materials and Methods

### Plant material and growth conditions

*A. filiculoides*, collected from a small pond of the botanical garden of the University of Perugia–Italy and characterized as reported in Costarelli *et al*. (2021), was grown in a climatic chamber under a 14 h light photoperiod at an irradiance of 220 μmol m^−2^ s^−1^ (photosynthetically active radiation - PAR) and a light/dark temperature of 25/19°C. The solution employed for *A. filiculoides* stock maintenance was the Watanabe’s growth solution (Watanabe *et al*., 1992), which was completely replaced every 7 days and refilled when necessary. Once a week*, A. filiculoides* stock cultures were split to avoid overcrowding and gently disentangled to favour growth. After removing the hull with tweezers, the seeds of rice *Oryza sativa* L. var. Kitaake were surface sterilized with ethanol for 30’’, followed by 30’ in 30 mL of commercial bleach and 2 μL of Tween20 in falcon tubes under continuous shaking. Afterward, 8-10 washes with sterile water were performed under a laminar flow hood. Seeds were left in imbibition in the dark o/n. The seeds were sowed in Petri dishes containing half-strength MS nutrients (½MS) (Murashige and Skoog, 1962) for 2 days in the dark at 30°C for germination, and successively in Magenta boxes for 10 days for seedlings growth. Four 13-day-old seedlings per replicate were then transplanted into specifically designed plex hydroponic boxes (230×230×150 mm) harbouring 4 pivots sticking out of the base and containing ca. 5 L of Yoshida’s nutritive solution (Gregorio *et al*., 1997). Each pivot could hold a hydroponics net pot that, once filled with expanded clay taken in place by a piece of non-woven tissue, supported rice growth for the entire duration of the experiment. Boxes containing sole rice plants (control, hereinafter referred to as R treatment) or rice co-cultivated with *A. filiculoides* (hereinafter referred to as R+Af treatment) were kept in a growth chamber under the conditions described above for *A. filiculoides* maintenance. The surface of the boxes without Azolla were covered and the walls of all the boxes wrapped with aluminium foil to prevent the development of algae. The growth solution was completely replaced every 2 weeks and refilled when necessary. No water-aeration system was applied and, when necessary, dying *A. filiculoides* plants substituted with fresh ones to ensure the same level of Azolla coverage. Unless differently specified, at least three boxes per treatment with four rice plants each were employed as replicates for each experiment and the entire experimental set up was run three times.

### Morphological analysis of rice plantlets

Number of leaves and tillers and the height of rice plants grown under R+Af and R treatments were scored every week for 63 days from the start of the hydroponics cultivation, herein after referred to as Days After Transplanting (DAT). Root architecture and fresh/dry plant weight were monitored at 15, 30, and 60 DAT by harvesting 1 plant for each replicate. The harvested plants were dipped in water to remove the adhering clay particles and other macroscopic contaminants and then carefully dried. The length of adventitious (ARs) and lateral (LRs) roots and that of shoots were measured using a ruler. Roots were then detached from the stems to measure the fresh weight of below and above water surface organs. The number of ARs and tillers were counted in all plants, whereas the number of ARs with and without LRs were counted in a subset of 3 plants per treatment. Portions of ARs, collected at 150 mm from the base of the stem, were observed at the epifluorescence light microscope (excitation 495 nm; emission 515 nm; long-pass filter of 515 nm) to determine the frequency of LRs (expressed as number/mm). The dry weight of root and leaves was measured by treating a portion of these organs in the oven at 80°C until a constant weight was reached.

### Determination of inorganic nitrogen content in culture media

The concentration of different forms of inorganic nitrogen in the media from R and R+Af treatments was evaluated at 10, 20 and 30 DAT. NH_4_^+^ concentration was assessed by the spectrophotometric absorbance at 690 nm of a derivative of indophenol formed by reaction of sodium salicylate and chlorine ammonia in an alkaline environment as reported in Jeong and colleagues (2013). The absorbance at 420 nm of the products resulting from the reaction between nitrates and sodium salicylate in acid solution for sulfuric acid was employed to assess the levels of N0_3_^-^ (Monteiro *et al*., 2003). NO^2-^ was determined following Monteiro *et al*. (2003): at pH 2.0-2.5 the sulfanilamide (I) is diazotized by nitrous acid and the resulting diazo compound is coupled with N-(1-naphthyl)-ethylenediamine (II) to form an azocompound that absorb at 543 nm (Monteiro *et al*., 2003; Jeong *et al*., 2013).

### Nitric oxide quantification in rice roots

Nitric Oxide (NO) was analysed in the AR apex cells of rice grown under R+Af and R treatments at 15 and 30 DAT. NO yellow–green fluorescence was detected by epifluorescence light microscopy (excitation 495 nm; emission 515 nm; long-pass filter of 515 nm) after incubation of a root portion in the dark in 10 μM 4,5-diaminofluorescein diacetate (DAF-2DA) for 3 h at RT (Kojima *et al*., 1998). DAF-2DA permeates through the cell membrane and was hydrolysed to DAF-2, which was retained in the cell owing to its relatively poor permeability. DAF-2 reacts with NO to form fluorescent DAF-2 T. NO content in the roots of rice grown with and without *A. filiculoides* were computed using ImageJ v. 1.4 and the corrected total cell fluorescence (CTCF) was calculated as: CTCF = Integrated Density − (Area of selected cell × Mean fluorescence of background readings) NO contents were also evaluated via the colorimetric Griess assay according to manufacturer instructions (Sigma-Aldrich). The oxidation product of NO, nitrite, reacts with sulfanilamide and N-(1-naphthyl) ethylenediamine dihydrochloride (NED) to yield a pink stable azo product. The conversion of spectrophotometric absorbance values recorded at 548 nm to nitrite concentration, and hence indirectly NO, was possible using the linear regression of the calibration curve (Vishwakarma *et al*., 2019). The calibration curve was done according to the manufacturer instructions and NO content was calculated by comparison with a standard curve of NaNO_2_ as described in Zhou *et al*. (2005).

### Rice root transcriptomics analysis

To collect samples for untargeted RNA analysis, rice plants from both treatments were rinsed with distilled water and the distal portions, up to 1 cm from the apices, of rice roots were cut with a sterile blade and immediately frozen in liquid nitrogen. If necessary, samples from different plants were pooled. RNA from the roots of three biological replicates per treatment at 15 DAT was isolated with Qiagen RNeasy Plant Mini Kit and treated with Qiagen Rnase-Free Dnase Set. Stranded mRNA libraries were prepared by Novogene with poly-T oligo-attached magnetic beads, followed by random hexamer priming and second strand cDNA synthesis with dUTP. The libraries were sequenced in paired-end 150bp mode on Illumina platform. To remove adapters, incomplete reads and low-quality reads, raw reads were filtered and quality-trimmed with Fastq-mcf (Aronesty, 2011) with options -l 50 -q 30 and quality of read sets was assessed with FastQC (Andrews, 2021) and MultiQC (Ewels *et al*., 2016). Reads were mapped on Oryza sativa cv. Nipponbare Os-Nipponbare-Reference-IRGSP-1.0 with annotation version 2022-03-11 (downloaded from the RAP-DB database, available at rapdb.dna.affrc.go.jp/download/irgsp1.html) (Kawahara *et al*., 2013; Sakai *et al*., 2013). For mapping, reads alignment and transcript quantification were performed with Rsem v1.3.3 (Li and Dewey, 2011). Count tables were imported in R v4.0.2 with package TxImport v1.16.1 (Soneson *et al*., 2015) and differential gene expression analysis between growth conditions was performed with DeSeq2 v1.28.1 (Love, Huber and Anders, 2014). Finally, the list of differentially expressed (DE) genes with criteria |logFC|> 0 and padj ≤ 0.05 (unless specified otherwise) was selected to run functional enrichment analysis on ShinyGO v0.76 (Ge *et al*., 2020), using the set of terms from Gene Ontology (Biological project, Molecular Function and Cellular Component) and KEGG, and using the list of all genes with detectable expression as background. ShinyGO was also used to investigate differentially expressed genes (DEGs) and produce a network summarizing enriched pathways. Pathways were linked when they shared at least 20% of genes. The resulting network was visualized on CytoScape v3.9.1 (Shannon *et al*., 2003).

### Validation of RNA-seq data by qRT-PCR

To validate the dataset obtained by RNA-seq analysis, the expression of a subset of DEGs was monitored by quantitative reverse transcriptase PCR (qRT-PCR) analysis. In Table S1 are given the 12 target genes and the 4 housekeeping genes amplified along with their primer pairs. Total root RNA was isolated from an independent experimental set of rice samples compared to those employed for RNA-seq analysis. Root samples, collected as reported in the previous paragraph, were taken at 0, 5, 9, 12 and 15 DAT. One and half µg of RNA isolated as reported above was reverse-transcribed in the presence of Maxima H Minus Reverse Transcriptase (Thermo Fisher Scientific, Milan, Italy) and 100 pmol of random hexamers (Euroclone, Milan, Italy), in a final volume of 20 µL. An aliquot of 2 µL of 1:10 diluted cDNA was used in the PCR reaction, which was carried out using the BlasTaq 2X qPCR Mater Mix (ABM, Richmond, Canada) in an Light Cycler 86 apparatus (Roche) using the following cycling parameters: an initial step at 95°C for 60’’, 30 cycles of a step at 95°C for 10’’ and a step at 60°C for 30’’, followed by a high resolution melting curve performed as: 95°C for 60’’, 40°C for 60’’; 65°C for 1’’ and 97°C for 1’’, prior to the cooling step at 37°C for 60’’. For each gene, time point and biological samples four technical replicates were amplified. For each primer pair, the efficiency of PCR was tested as reported previously (Escaray *et al*., 2014). The 2^−ΔCt^ gene expression quantification method was applied to compare the relative expression levels among the target genes (Bizzarri et al. 2020). Here, the 2^-ΔCt^ method was based on the differences between the relative expression levels of the target genes and the geometric mean of the 4 housekeeping genes described in de Castro Dos Santos *et al*. (2018).

### LCMS analysis of phytohormones in rice roots, leaves and growth media

Extraction and purification of phytohormones from plant material (i.e., rice roots and leaves) and growing media at 0 and 15 DAT were performed by solid-phase extraction (SPE) followed by LCMS-QQQ according to a previously published protocol with minor modifications (Prerostova *et al*., 2021). Root sampling was performed as reported for RNA-seq and qRT-PCR analyses. For leaf sampling, the median part of the most expanded leaves was collected. For the media, 1 ml aliquots were collected for each box and then lyophilized. Tissue disruption (30mg) was performed in 100 μL extraction solvent (1M HCOOH) using a benchtop homogenizer FastPrep-24 (MP Biomedicals, CA, USA) in an extraction solvent and about 0.05 g of 1.5 mm zirconiumoxide balls. This step was not performed on media samples, which were resuspended in 100 μL of extraction solvent. Ten μL of the internal standards (IS) were added to the samples, thoroughly mixed and centrifuged at 4°C and 30,000*g.* The resultant supernatants were gently collected and applied to the SPE plate. The following isotope-labelled mixture was added to each sample: ^13^C_6_-IAA (Cambridge Isotope Laboratories, Tewksbury, MA, USA); ^2^H_4_-SA (Sigma-Aldrich, St. Louis, MO, USA); ^2^H_3_-PA, ^2^H_3_-DPA (NRC-PBI); ^2^H_6_-ABA, ^2^H_5_-JA, ^2^H_5_-tZ, ^2^H_5_-tZR, ^2^H_5_-tZRMP, ^2^H_5_-tZ7G, ^2^H_5_-tZ9G, ^2^H_5_-tZOG, ^2^H_5_-tZROG, ^15^N_4_-cZ, ^2^H_3_-DZ, ^2^H_3_-DZR, ^2^H_3_-DZ9G, ^2^H_3_-DZRMP, ^2^H_7_-DZOG, ^2^H_6_-iP, ^2^H_6_-iPR, ^2^H_6_-iP7G, ^2^H_6_-iP9G, ^2^H_6_-iPRMP, ^2^H_2_-GA_1_, ^2^H_2_-GA_4_, ^2^H_2_-GA_8_, ^2^H2-GA_12_, ^2^H_2_-GA_19_, ^2^H2-GA_34_, (^2^H_5_)(^15^N_1_)-IAA-Asp, (^2^H_5_)(^15^N_1_)-IAA-Glu (Olchemim, Olomouc, Czech Republic), (^2^H_5_)(^15^N_1_)-IAM, ^2^H_5_-IAA-GE+Am, ^2^H_4_-OxIAA, ^2^H_4_-OxIAA-GE+Am and ^2^H_3_-epi-Br.

The 96-well SPE plates (Oasis HLB 10 mg sorbent per well; Waters, Milford, MA, USA) were activated by sequentially applying 100 μL of acetonitrile, water and extraction solvent for 3 minutes. Pressure was applied to pass the samples through the extraction columns using a Pressure+96 positive pressure manifold (Biotage, Uppsala, Sweden). Pellets were re-extracted with an additional 100 µL of extraction solvent, centrifuged and applied again to the column plates. The wells were then washed 3 times with 100 µL of water. The phytohormones were eluted with 2 x 50 μL elution solvent (50% ACN in water). The eluate was collected in a collection plate, sealed with the 96-well silicone cap and stored at −20°C until LCMS analysis. Phytohormones were separated on Kinetex EVO C18 column (2.6 μm, 150 x 2.1 mm, Phenomenex, Torrance, CA, USA). Mobile phases consisted of A— 5mM ammonium acetate and 2 μM medronic acid in water and B—95:5 acetonitrile:water (*v/v*). The following gradient was applied: 5% B in 0 min, 5–7% B (0.1–5 min), 10–35% B (5.1–12 min) and 35–100% B (12–13 min), followed by a 1 min hold at 100% B (13–14 min) and return to 5% B. Hormone analysis was performed using an LCMS system consisting of a UHPLC 1290 Infinity II (Agilent, Santa Clara, CA, USA) coupled to a 6495 Triple Quadrupole Mass Spectrometer (Agilent, Santa Clara, CA, USA) operating in MRM mode, with quantification by the isotope dilution method. Data acquisition and processing were performed using Mass Hunter software B.08 (Agilent, Santa Clara, CA, USA). The amounts of the quantified compounds were expressed as pmol/gFW and pmol/mL. Outliers were identified by statistical test (i.e., interquartile range IQR) and excluded for subsequent analysis in MetaboAnalyst (version 5.0) (Pang *et al.*, 2021).

### *T. azollae* growing conditions and phytohormone analysis of the medium

The cyanobacterium *T. azollae* used in this study was obtained from the Culture Collection belonging to the Institute of BioEconomy (IBE), National Research Council of Italy (CNR; Sesto Fiorentino, Firenze, Italy). The cyanobacterium was cultivated in laboratory in batch mode using vertical glass column reactors (5 cm light path, 600 ml working volume) and BG110 (nitrogen-free) as culture medium (Rippka *et al*., 1979). Continuous illumination of 40 μmol m^-2^ s^-1^ (PAR) was provided by means of cool white lamps (Dulux L, 55W/840, Osram), and a culture temperature of 22 ± 2°C was maintained by thermostat-cultivation room. Cultures were bubbled with a sterile air/CO_2_ mixture (98/2, v/v) to ensure continuous mixing, remove dissolved oxygen, and maintain pH within the desired range (7.5–8.0). The cultures were diluted once a week by repeating the same growth cycle three times (i.e., 200 mL of culture was removed and processed, and the same volume of fresh culture medium added). For phytohormone analysis of the *T. azollae* medium, samples were taken in sterile conditions, centrifuged at 4,000 *g* for 25’, the supernatant filtered (pore size 0.45 µm), aliquoted into vials and stored at −20°C until analysis.

### Statistical analysis

Rice plants were grown with and without *A. filiculoides* and experiments were repeated thrice. For each treatment, 12 to 16 plants were independently computed unless specified otherwise. To determine significant differences between treatments, data were tested for homogeneity of variance (F-test or Levene’s test) and normality distribution (Shapiro-Wilk’s test). If the assumptions of these tests were not violated, data were analyzed via unpaired two-sample t-test or via one-way analysis of variance (ANOVA) and post hoc comparison (Tukey’s HSD), according to the number of treatments. If the assumptions of the homogeneity of variance or normality distribution tests were violated, data were analyzed via the Wilcoxon rank sum test. Statistical analyses were carried out in R studio (version 3.5.3) (R Core Team, 2016) or in MetaboAnalyst (version 5.0). The level of significance was set to P < 0.05 or to FDR cut-off < 0.05 and treatment mean values ± S.E. were plotted unless differently specified. To analyze targeted-harmonics data, multivariate statistics allowed for the computation of sPLSD and respective Loadings plot after data normalization by median and Pareto scaling (i.e., mean-centered and divided by the square root of the standard deviation of each variable). Statistical analysis was performed both at the same time point between treatments and over time within the same treatment.

## RESULTS

### The co-cultivation with *A. filiculoides* changed rice root architecture and induced a NO boost in the rice adventitious roots

To investigate the effects of *A. filiculoides* on rice RSA, the number and length of ARs, LRs, and the root weight in the R+Af and R treatments were measured at 15, 30, and 60 DAT. A significant increase in the number of ARs was seen in the presence of *A. filiculoides* (Fig. 1A) at 30 DAT. At the same time point, the length of the ARs in the R+Af treatment was shorter (Fig 1B), whereas at 60 DAT, for this trait, an opposite pattern was observed (Fig. 1B). The number of ARs provided with and without LRs, the length of LRs and the number of LR/mm were also evaluated in R and R+Af treatments at 15 and 30 DAT. Conversely, the monitoring of these parameters at 60 DAT was prevented by the extensive root mats adhering to the clay and the pot in all plants. At 15 DAT, the number of ARs provided with and without LRs was not significantly different between the treatments (Fig. 1C). Rather, at 30 DAT the number of ARs provided with LRs was significantly higher in the R+Af compared to the R treatments (Fig. 1D). Differently, the number of thick, unbranched ARs and the ratio of the number of ARs having LRs divided by that without LR (AR +LR/-LR) were not significantly affected. The length (μm) and the number/mm of rice LRs were neither significantly different at 15 nor 30 DAT (data not shown). At 15, 30 and 60 DAT no significant differences between the two treatments were observed for the root weight. However, in the presence of *A. filiculoides* the root biomass increased of 32% and 48% at 30 and 60 DAT, respectively. It is worth noticing that the P value of the comparison between treatments at 60 DAT was P = 0.06 (Fig. S1).

**Figure 1.**
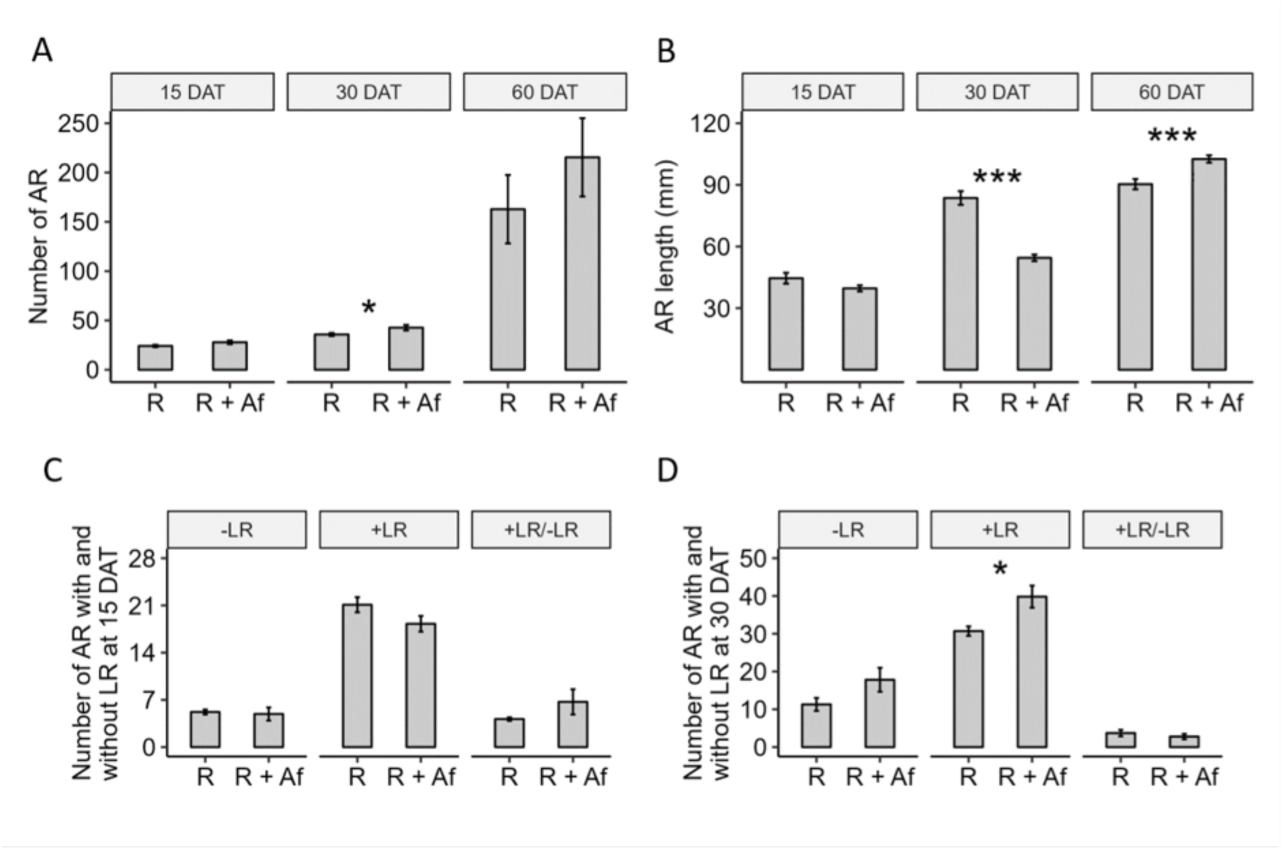
Effect of *A. filiculoides* on the number (A) and length (B) of ARs in rice at 15, 30 and 60 DAT, and on the number of LRs of rice at 15 (C) and 30 DAT (D). Bar Graphs represent mean values ± SE of control rice (R) and rice grown with A. filiculoides (R+Af). A: 15 DAT R n = 6; 15 DAT R + Af n = 7; 30 DAT R and R+Af n = 18; 60 DAT R n = 7; 60 DAT R + Af n = 9. B: 15 DAT R n = 144; 15 DAT R + Af n = 187; 30 DAT R n = 254; 30 DAT R + Af n = 333; 60 DAT R n = 853; 60 DAT R + Af n = 1711. C and D: 15 and 30 DAT, respectively. R n = 10; R + Af n = 11. ***, P < 0.001. *, P < 0.05.

To gain a preliminary insight into the physiological events underlying the observed effects of *A. filiculoides* on rice RSA, the relative content of NO in the ARs from R and R+Af treatments was evaluated at 15 and 30 DAT by fluorescence analysis, and the Griess assay. Both analyses showed a significant increase in NO content in the ARs of rice grown in the presence of *A. filiculoides* at 15 DAT (Fig. 2; Fig S2).

**Figure 2.**
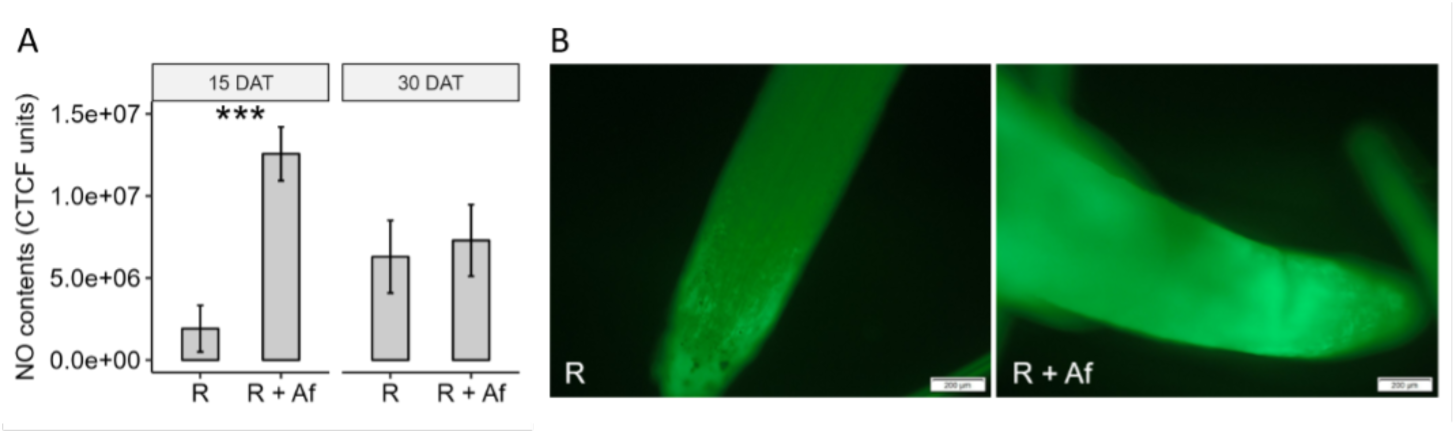
Effect of *A. filiculoides* on the relative amount of NO in the ARs of rice at 15 and 30 DAT. Bar Graphs (A) represent mean values ± SE of control rice (R) and rice grown with A. filiculoides (R + Af). 15 DAT R n = 23; 15 DAT R+Af n = 22; 30 DAT R and R + Af n = 15; 60 DAT R n = 21. ***, P < 0.001. CTCF: Corrected Total Cell Fluorescence. Pictures (B) were taken at the fluorescence microscope, and picture analysis was performed in ImageJ. Data shown do not take into account sample autofluorescence.

### *A. filiculoides* improved vegetative growth of rice plants and modified carbon allocation over time

The fern affected the growth of rice above-ground organs over time (Fig. 3). A significant increase in the number of leaves (Fig. 3A), plant height (Fig. 3B), and number of tillers (Fig. 3C) in the R+Af vs R treatment started to appear at 28, 14 and 28 DAT, respectively. However, the significance in the number of tillers was lost after 60 DAT. The biomass of rice grown under the two treatments was measured at 15, 30 and 60 DAT (Fig. S3). The aerial part and total rice weights tended to be slightly lower in the R+Af compared to the R treatment at 15 DAT, whereas at 30 DAT they significantly increased at 62.5% and 54%, respectively, and total plant fresh weight became higher in the R+Af than in R treatment (Fig. S3B). The increase in rice total biomass in the R+Af treatment at 30 DAT primarily resulted from that of the aerial part rather than from the roots (Fig. S1). At 60 DAT, the above-ground part and total rice weight again tended to be higher in the R+Af compared to the R treatment (P = 0.057; 58 and 74% increase, respectively). To assess whether *A. filiculoides* significantly affected rice carbon allocation over time, the root/shoot ratio of the fresh weights (FWs) at 15, 30, and 60 DAT were computed (Fig. 3D). A statistically significant decline was observed in the R+Af treatment vs R treatment at 30 DAT. At 60 DAT, the root-to-shoot ratio showed an opposite trend, as it tended to be higher in the R+Af treatment. Thus, co-cultivation with *A. filiculoides* affected carbon allocation in rice at 30 DAT, that is when rice in the R+Af treatment allocated significantly more carbon towards the aerial part compared to the root system in the R+Af treatment.

**Figure 3.**
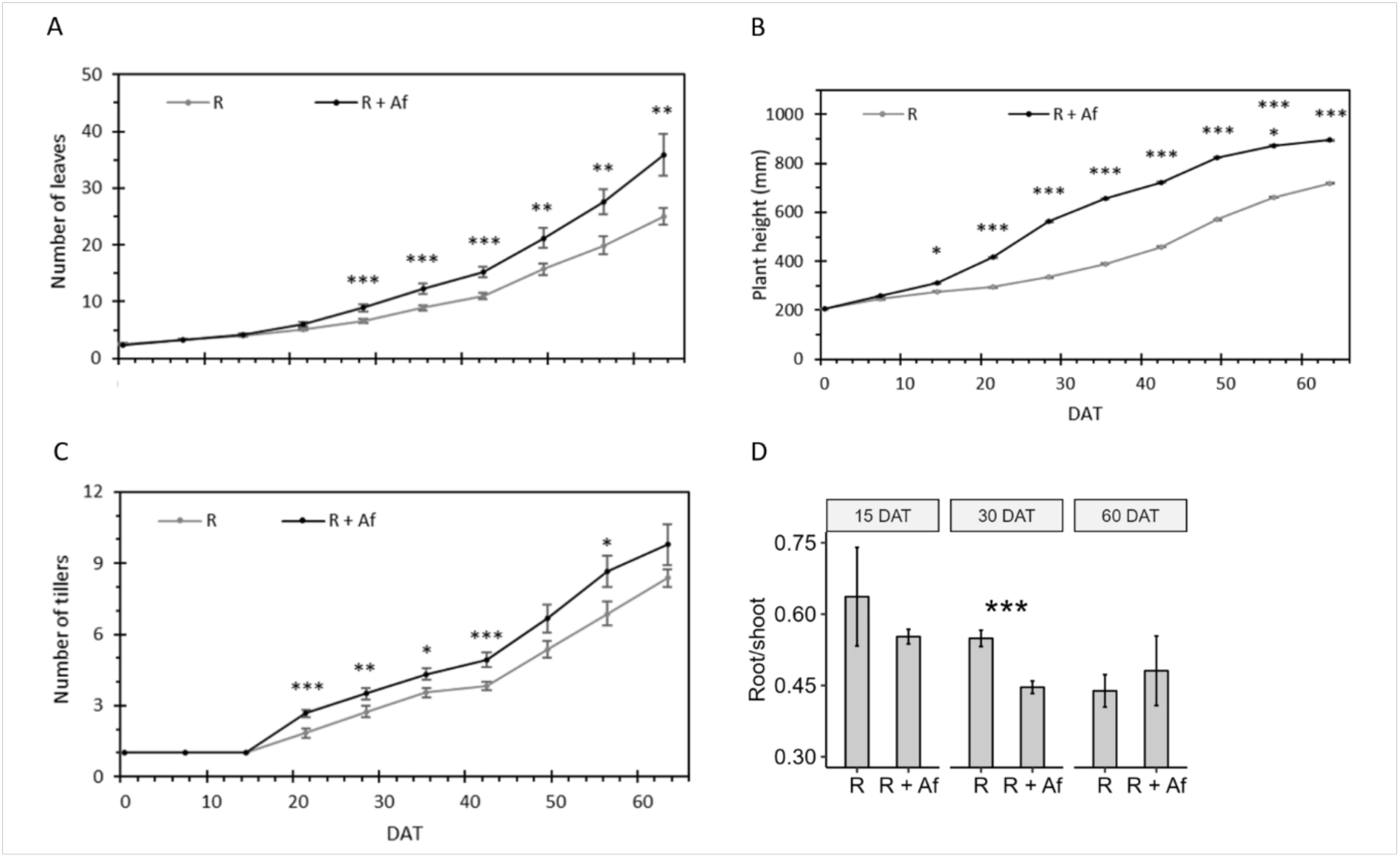
Effect of *A. filiculoide*s on the number of leaves (A), plant height (B), number of tillers (C) and root/shoot fresh weights (D) over two months. Linear Graphs represent mean values ± SE of control rice (R) and rice grown with *A. filiculoides* (R+Af). Linear graphs: 0, 7, 15, 20, 28 DAT R and R+Af n= 12; 34 and 39 DAT R n= 11; 34 and 39 DAT R + Af n = 12; 46, 53 and 60 DAT R n= 8; 46, 53 and 60 DAT R+Af n=9. Bar graphs: 15 DAT R n=6; 15 DAT R + Af n=7; 30 DAT R and R + Af n=20; 60 DAT R n=12; 60 DAT R + Af n=15 *, P < 0.05; **, P < 0.01; ***, P < 0.001.

### The inorganic nitrogen content did not increase in the media of co-cultivated rice

To investigate whether the changes in morphology and developmental timing of rice organs upon co-cultivation with *A. filiculoides* were related to the increase of inorganic N in the growth media, the levels of inorganic N forms were assessed at 3 time points, namely 10, 20 and 30 DAT. At any time point, the levels of inorganic N increased in the media where *A. filiculoides* was present (Table S2). Rather, some inorganic forms decreased in the media with *A.filiculoides,* although only at the first time points.

### *A. filiculoides* altered rice root transcriptome at 15 DAT

Morphological changes triggered by *A. filiculoides* in rice plants occurred at 30 DAT for the roots and a few days earlier for the aerial parts. By reasoning that these changes resulted from signals exchanged at root levels between the fern and rice plants since the early stage of co-cultivation, root transcriptomics using control and *A. filiculoides* co-cultivated rice plants was assessed at 15 DAT via RNA-seq analysis. The similarity between the gene expression profiles of the replicates of the same treatment was assessed via PCA (FDR cut-off < 0.05) (Fig.4A). The dimensional-reduction analysis revealed that the three replicates of R treatment clustered together as much as did those of R+Af treatment. Among all annotated genes, 473 were differentially expressed (DE), of which 230 were upregulated and 243 were downregulated (FDR cut-off = 0.05) (Fig 4B). The functional enrichment analysis of the 473 DEGs at 15 DAT highlighted the 20 most enriched biological processes (BP) and molecular function (MF) terms, shown in Fig 5A and B, respectively (FDR cut-off = 0.001 for both). Notably, the methionine (Met) salvage pathway (MSP), the amino acids (aa) salvage pathway (ASP), and iron and metal transport and homeostasis were the most enriched BP terms. As for MF, the terms iron and metal binding and transmembrane transporters were the most enriched. No results were obtained selecting the pathway database ‘GO cellular component’ (CC). Among the 230 upregulated DE genes, the most enriched BP was intracellular sequestering of iron ion, whereas the response to reactive oxygen species (ROS) was the BP with the highest number of genes (FDR cut-off = 0.001). The most enriched MFs were ferric and ferrous binding (FDR cut-off = 0.001) (Fig. S4A and B). The same analysis of the 243 downregulated DE genes revealed that the most enriched BPs were L-Met recycling and iron and metal ions transport (FDR cut-off = 0.001), and the most enriched MF was metal ions transport (FDR cut-off = 0.001) (Fig. S5 A and B).

**Figure 4.**
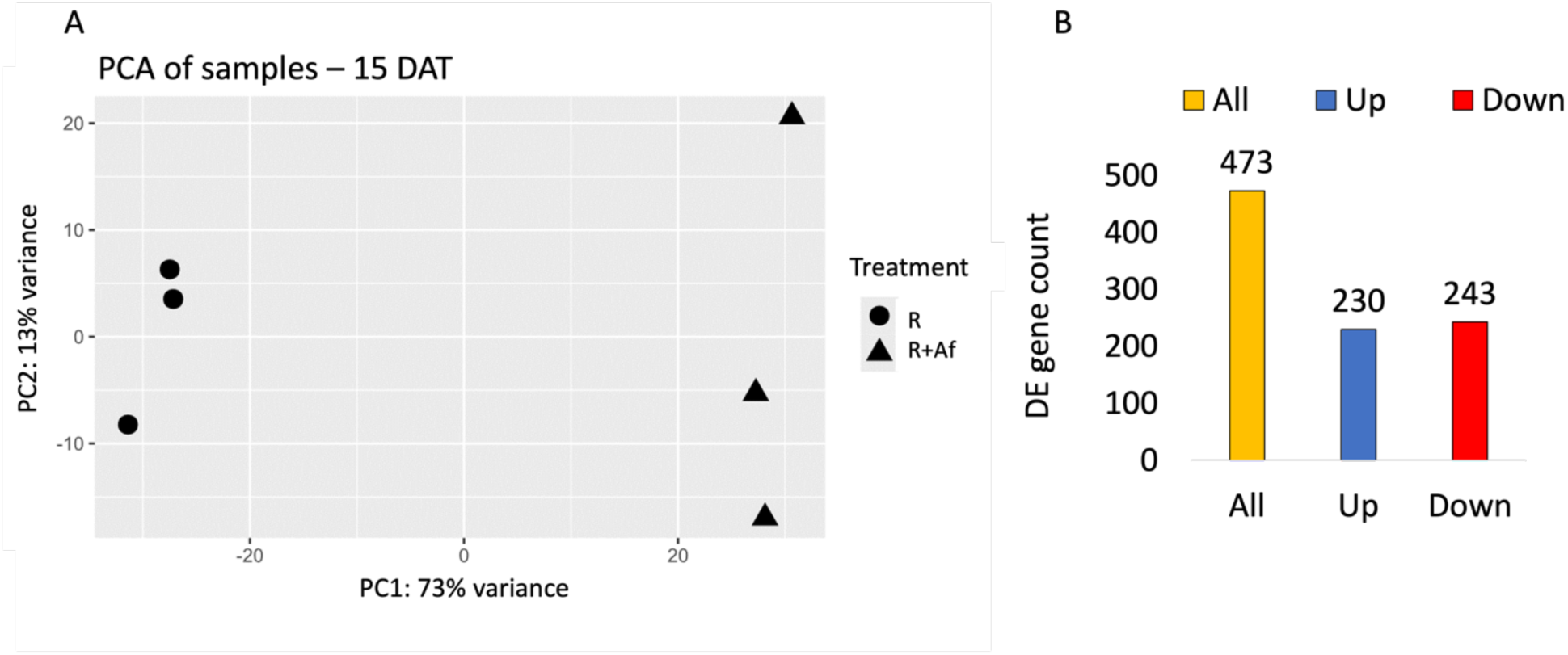
PCA (A) and DE gene count at 15 DAT (B).

**Figure 5.**
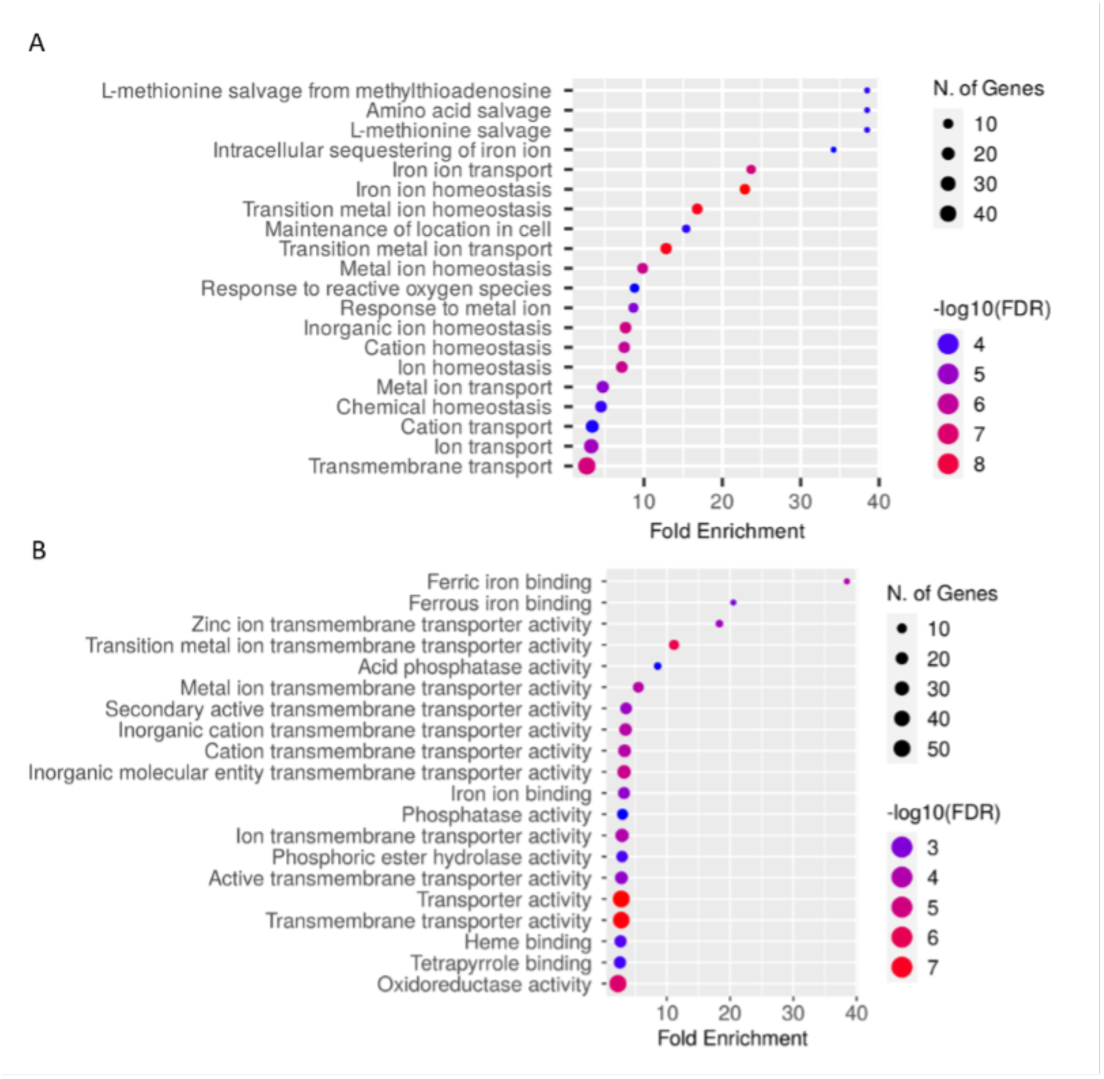
The 20 most enriched BP (A) and MF (B) terms among the DEGs in rice roots at 15 DAT.

The MSP is part of the cysteine and methionine metabolism, which also comprises the biosynthesis of cysteine (M00021) and ethylene biosynthesis (M99368). Therefore, the MSP was further investigated. All the DEGs involved in the pathway were downregulated (Table S3). Among the most DE genes, those related to iron homeostasis were also included (Table S4). Rice employs two strategies for iron solubilization and uptake (Ishimaru *et al*., 2006, Li *et al.*, 2023), and genes related to both strategies were severely downregulated, whereas those for iron storage in plastids (*Ferritin 1*, *FER1, FC=2.45* and *ferritin 2*, *FER2, FC=2.49*) and vacuoles (*Vacuolar Iron Transporter 1*, *VIT1-2*, FC=6.07 and the *Vacuolar Iron Transporter Homolog 2*, *VITH2*, FC= 6.44) upregulated in the R+Af treatment (Fig.6, Table S4).

**Figure 6.**
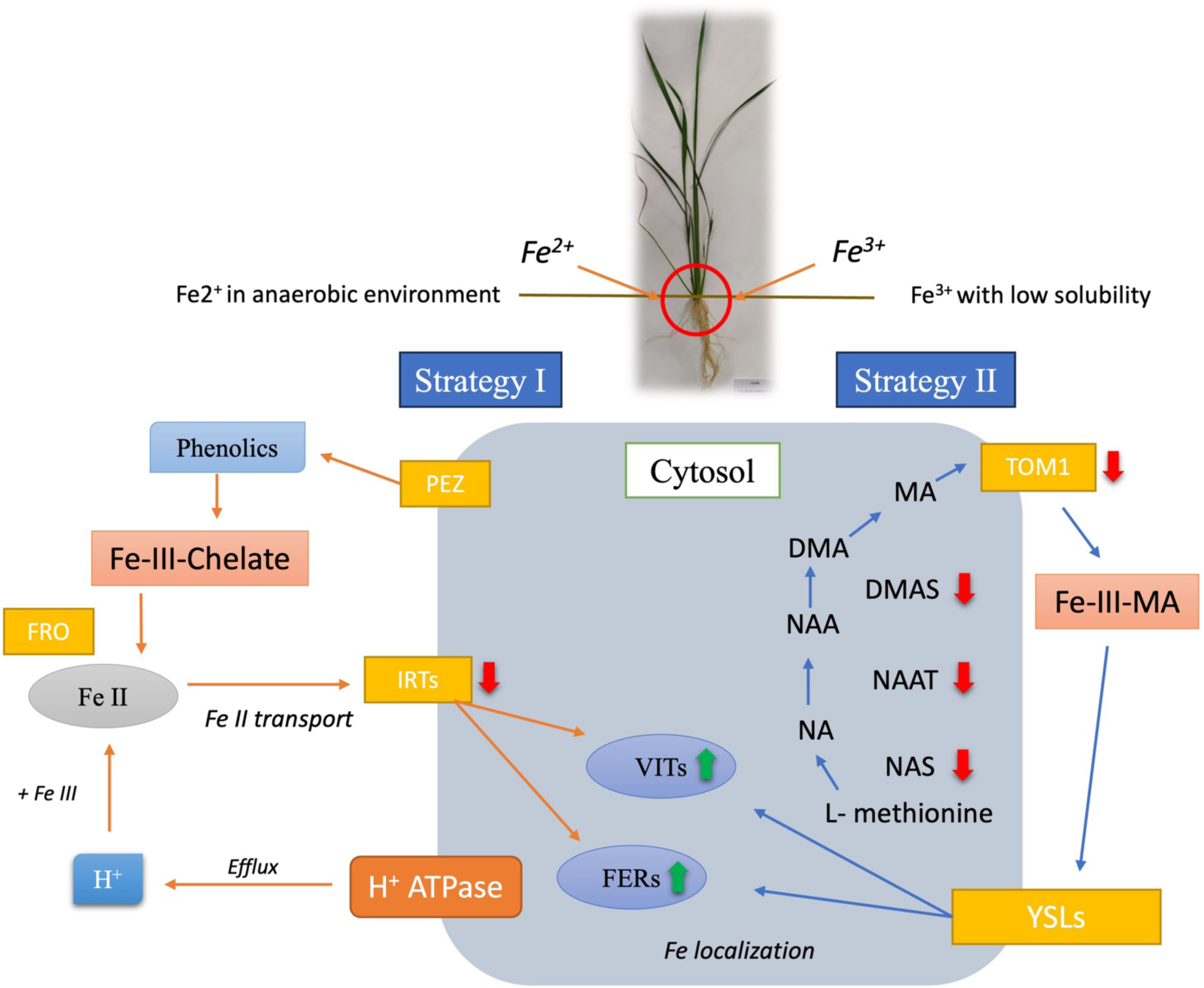
Schematic representation of the iron acquisition systems in rice roots. Under Strategy I the ferric chelate complex is reduced to ferrous ion by FRO. H^+^ released by the proton pump acidifies the medium to convert ferric ion to ferrous ion. IRTs transport the ferrous ion into the root cells. VITs are responsible for transportation and accumulation of iron within the vacuole while FERs are the proteins accommodating excess iron within the cells. Under Strategy II, MA phytosiderophores, produced by the sequential activity of NAS, NAAT and DMA, chelate Fe(III). Chelated Fe(III) complex is transported across the membrane into the root cell by YSLs. Green and red arrows refer to transporters/enzymes whose mRNA are increased and decreased, respectively, in the R+Af vs R treatment. For the differential expression of regulatory genes refer to TableS4. The picture was redrawn from Kar and Panda (2020) and Kobayashi *et al*., (2014).

To investigate the link between iron homeostasis and methionine metabolism, a network analysis was performed by querying all DEGs (Fig. 7). The functional categories were linked if they shared ≥ 20% genes (FDR cut-off = 0.05). The results showed that the nodes – ‘L-Met salvage pathway (MSP)’ and ‘iron homeostasis’ – were connected to the node ‘response to NO,’ and are involved in response to stimulus, transport, and homeostasis of ions, in particular iron, and amino acid metabolism. Intriguingly, the terms L-Met salvage pathway, iron homeostasis, and response to NO shared a common gene, *Iron deficiency-induced protein 2 (IDI2)*, which was downregulated (FC= −3.32) in the presence of *A. filiculoides* (Table 1 and S3). Within the 15 DAT dataset, three NO-related genes were found to be upregulated: the *Nitrate reductase 2* (*NR2*), which encodes the NO biosynthetic enzyme, and two *Nitrate transporter (NRT)* genes (Table 1).

**Figure 7.**
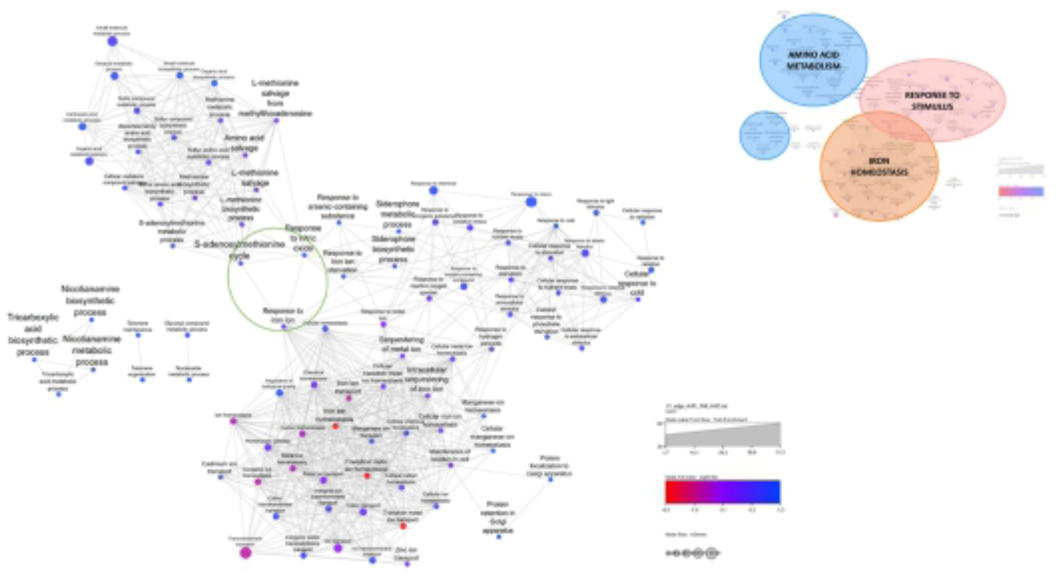
Network analysis of DEGs. The functional categories are linked if they shared ≥ 20% genes (FDR cut-off = 0.05).

**Table 1.** NO-related DEGs at 15 DAT.

| RAP-DB | Symbol | Log2FC | padj | Description |
| --- | --- | --- | --- | --- |
| *Os02g0770800* | *NR2/NIA1/NIA2* | 5.373 | 8.4E-05 | *NADH/NADPH-dependent nitrate reductase* |
| *Os10g0554200* | *NRT1.1B/NPF6.5* | 3.049 | 3.9E-04 | *Nitrate transporter 1.1B* |
| *Os10g0169900* | *NRT* | 4.308 | 1.6E-04 | *Nitrate and chloride transporter* |

In the roots of *A. filiculoides* co-cultivated rice plants the higher levels of NO and *FER* mRNAs at 15 DAT were coupled with the upregulation of several genes coding for ROS scavengers such as *Catalase A1* (*CATA1,* FC=2.23), *Ascorbate peroxidase 1* (*APX1,* FC=1.60)*, Ascorbate peroxidase 7* (*APx7,* FC=.46), the *Glutaredoxin-like protein 4* (*GRL4*, FC=1.77) and the *gluthatione transferase 41* (*GSTU41,* FC= 1.77) (Table S5).

Since R+Af treatment induced a differential accumulation pattern of the bioactive forms of IAA, ABA, CKs and SA (see below), the dataset was analysed for phytohormone-related genes (Table S6). The identified DEGs were generally not connected to the biosynthesis of these phytohormones, rather to their signalling and response. In this respect, it is noteworthy the differential regulation of 4 auxin -(*Os01g0924966*, FC=6,43; *Os09g0545400*, FC=−1,14; *Os09g0133200*, FC=−1,02; *Os09g0133200*, FC=−1,02985), 3 ABA - (*Os02g0543000*, FC= 3,333083; *Os11g0167800*, FC= 1,109878; *Os02g0734600*, FC= 1,491478), along with 2 CKs - (*Os04g0442300*, FC=2,17; *Os12g0139400*, FC=2,33) responsive genes. In particular, *Indole-3-Butyric Acid response 1 (IBR1, Os09g0133200*) involved in the conversion of the auxin precursor IBA to active IAA (Frick and Styrader, 2018) was downregulated (FC=−1,02), whereas the *Gretchen Hagen 3.12* (*GH3.12*, *Os11g0186500*), belonging to a family that catalyzes the conjugation reactions of salicylic acid, jasmonic acid, and IAA with amino acids to control their homeostasis (Guo et al., 2022), was upregulated (FC=3.43). Two ethylene-responsive genes (*Os04g0549800*, FC=3,70; *Os02g0656600*, FC=1,36) were upregulated in rice roots from R+Af treatment. Conversely, SA-related genes were not differentially expressed (Table S6).

### Targeted qRT-PCR analysis validated and extended RNAseq results

To validate RNA-seq results at 15 DAT, and simultaneously investigate gene expression at earlier time points, the quantification of the expression of a subset of DEGs of interest was performed on rice roots from R+Af and R treatments by qRT-PCR (Table S1) sampled at 0, 5, 9, 12 and 15 DAT. Genes were selected according to their biological function and the expression levels in the RNA-seq dataset. Transcriptomics and qRT-PCR results were in good agreement. In fact, the significantly different expression of 10 out of the 12 genes tested by qRT-PCR at 15 DAT was confirmed. For the remaining 2 DEGs, the *basic Helix-Loop-Helix 58* (*bHLH58*) and the *Protein Phosphatase 2C* (*PP2C*), qRT-PCR analysis confirmed the higher levels of their mRNAs in the presence of *A. filiculoides*, although the increments in the expression were not significant (Fig. 8).

**Figure 8.**
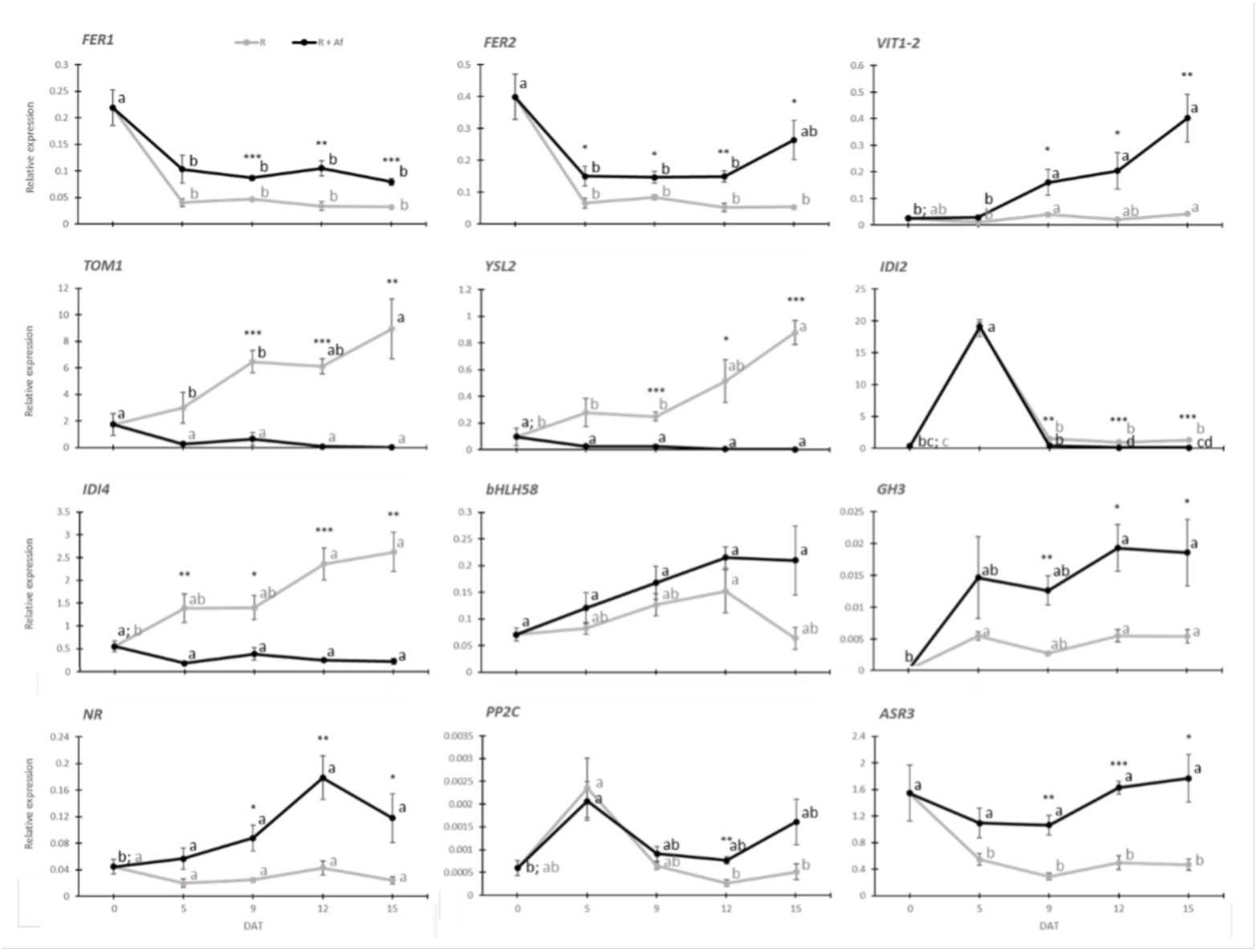
Genes expression profiles of selected DEGs at 15 DAT as resulted from the qRT-PCR analysis at 0, 5, 9 12 and 15 DAT. Asterisks indicate significant differences between the two treatments at a given time pint (t-test, P < 0.05), while letters indicate significant differences between time points of the same treatment (ANOVA, P < 0.05; Tukey’s HSD, P < 0.05).

Overall, the qRT-PCR analysis showed that the differential expression of the selected DEGs was evident much earlier than at 15 DAT. This occurred for the genes related to iron uptake/homeostasis (*FER1*, *FER2*, *VIT1-2*, *TOM1*, *YSL2*, *IDI2*, *IDI4*), with two of them, namely *FER2* and *IDI4*, differently expressed since 5 DAT, as well as for the gene related to NO biosynthesis (*NR*, since 9 DAT) and hormone maturation/signalling (*PP2C* at 12 DAT; *abscisic acid-stress- and ripening gene 3* -*ASR3,* since 9 DAT and *GH3* since 9 DAT).

### Targeted-hormonomics in rice organs and growth media in the presence and absence of *A. filiculoides*

To evaluate whether the morphological and transcriptional changes determined in rice by *A. filiculoides* could be related to an altered balance of phytohormones, the accumulation profiles of phytohormones, along with that of their precursors and metabolites, were evaluated in the roots and leaves of rice grown with and without *A. filiculoides* at 15 DAT. Furthermore, by assessing the phytohormonal contents in the growth media of rice grown under the two treatments, we aimed to gain insights into the molecular interplay occurring between *A. filiculoides* and rice. In Table S7, the hormonal compounds investigated are given.

### The presence of Azolla altered the levels of hormones in rice roots and leaves

Rice roots were sampled at 0 DAT (T_0_) before rice experienced the hydroponic cultivation and at 15 DAT under R+Af or R treatment. Discriminant Analysis (sPLSDA) showed a clear separation between roots from R and R+Af samples at 15 DAT (Fig. S6A). Additionally, the correspondent sPLSDA loadings plot highlighted the compounds that contributed most to explaining the differences between the treatments. The top ten compounds are reported in Fig. S6B and among them the most relevant was iP7G, followed by ABA.

At 15 DAT the levels of 5 out of the 48 quantified compounds were significantly different in the R + Af compared to the R treatment (Fig. 9, Table S8). These compounds belong to the cytokinin (CK), ABA, auxin and phenolic classes. The levels of three of them, namely ABA, the phytoalexin DPA, which is the product of irreversible ABA oxidation (Mongrand *et al*., 2003), and IAA-Glu were significantly higher in the R+Af treatment. Conversely, iP7G and the phytoalexin SinAc contents were significantly lower. At this time point, DZ9G was detected differently from 0 DAT, although its levels were not different between treatments. By filtering the data less stringently, the levels of two other compounds, cZROG and IAA-Asp, were significantly higher in R+Af at 15 DAT (unpaired two samples t-test, P < 0.05; Table S8).

**Figure 9.**
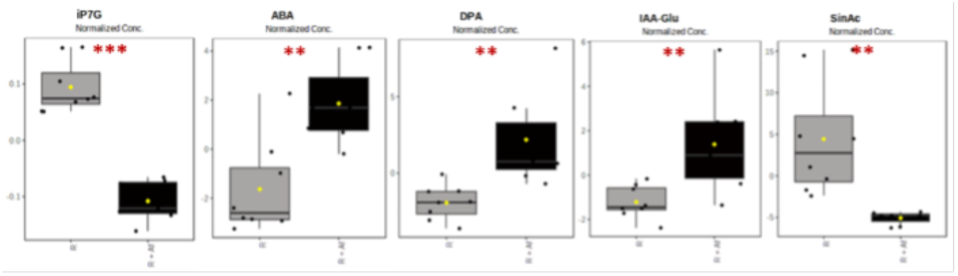
Differential levels of hormonal compounds in the roots of rice grown with and without *A. filiculoides* at 15 DAT. Unpaired two samples t-test for all five compounds except IAA-Glu, for which the unpaired two samples Wilcoxon test was performed. For all five compounds: P < 0.01, FDR cut-off ≤ 0.05. See also Table S8.

Looking at the root hormonal profiles from 0 to 15 DAT, it emerged that the presence of *A. filiculoides* changed the accumulation profiles of several bioactive and metabolic forms in rice roots. In Table S9 the levels of all compounds detected in the R and R+Af treatments are given, whereas Fig 10 shows the accumulation profiles of the bioactive forms. Among the bioactive forms, ABA levels increased from 0 to 15 DAT in both treatments, but their levels were significantly higher in the R+Af vs R treatment at 15 DAT (Fig. 10A).

**Figure 10.**
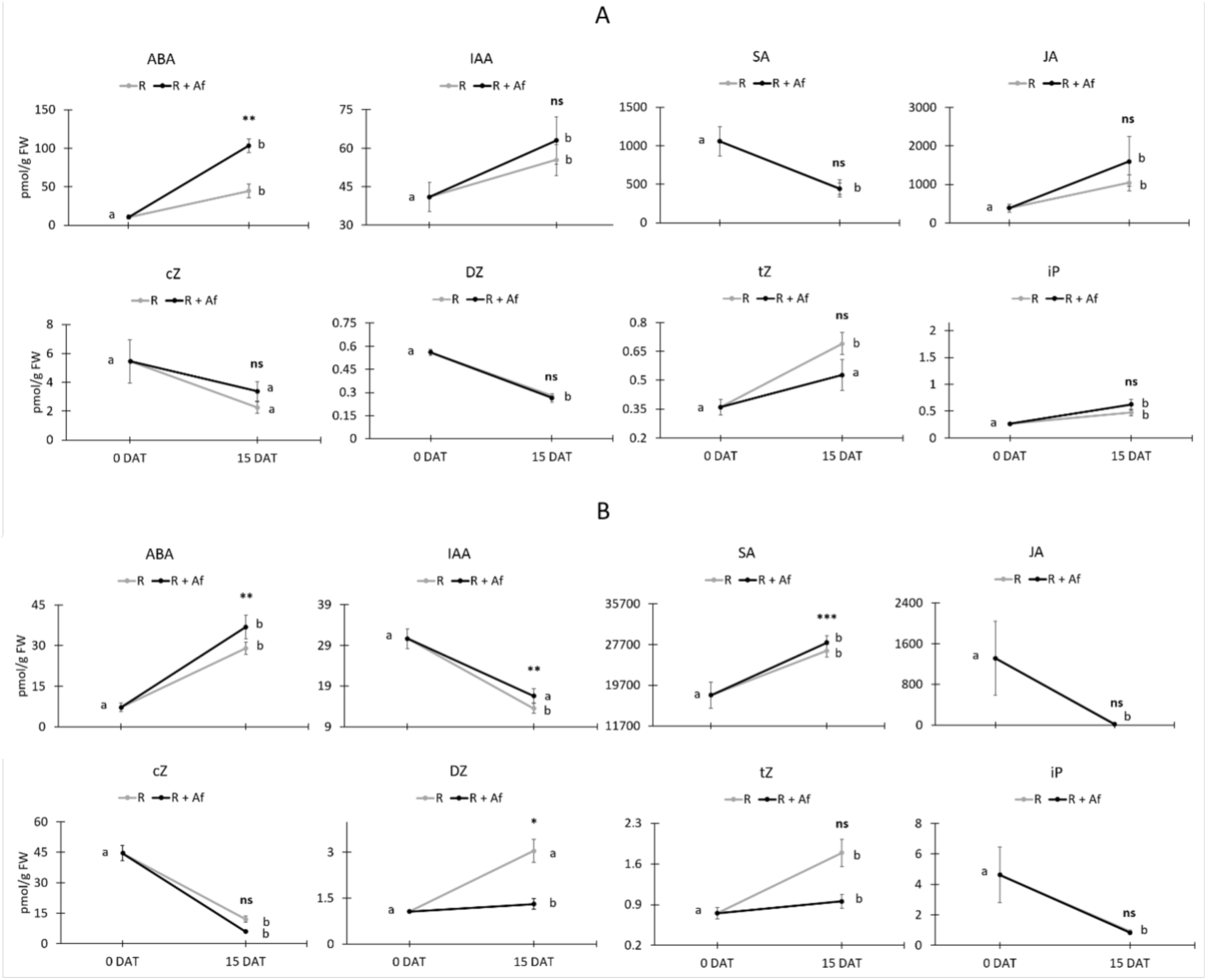
Accumulation profiles at 0 and 15 DAT of the bioactive hormones in rice roots (A) and leaves (B) from R+Af and R treatments.

Moving rice seedlings to hydroponic conditions caused a marked SA decrement and increased JA root contents, irrespective of treatment. A more complex picture emerged for the bioactive CKs: the levels of cZ did not decrease significantly at 15 DAT, whereas those of DZ did in both treatments. Conversely, the accumulation of tZ at 15 DAT increased slightly in the presence of *A. filiculoides*, but significantly in the control. Differently, iP levels increased significantly at 15 DAT, regardless of the treatment (Fig. 10A).

For what concerns leaves, the sPLSDA analysis displayed a marked separation of the hormonal compounds between R + Af and R treatments (Fig. S7A), with the oxidized forms of IAA, SA, and several CKs being the compounds that contributed the most to their separation (Fig. S7B). Of the 50 quantified compounds in rice leaves at 15 DAT, 19 significantly differed between the treatments (Fig. S8). In particular, among CKs, with the only exception for DZR, all the others (tZOG, DZOG, cZOG, cZROG, iPRMP) were accumulated at higher levels in the presence of *A. filiculoides* (Table S10). Under this condition, the 4 and 5 differentially accumulated ABA-related (ABA, ABA-GE, PA and 9OH-ABA) and IAA-related (IAA, IAA-Glu, IAM, I3A and ILacA) compounds, respectively, also showed higher levels. Likewise, the levels of 3 phenolic compounds (SA, BzA and PAAM) and the sole gibberellin detected (GA19) were higher in the presence of *A. filiculoides* (Fig. S8). By filtering the data less strictly, the accumulation of 10 additional compounds was significantly different. These were tZRMP, DZ, cZRMP, with lower levels and iP7G, iP9G, MeS-ZR, MeS-iP, DPA, JA-Me and OxIAA-GE with higher levels (unpaired two samples t-test; P < 0.05; Table S10) in the R+Af vs R treatment. By looking at the hormonal levels over time, it emerged that the presence of *A. filiculoides* changed those of several bioactive (Fig 10B) and metabolic forms (Table S11) in rice leaves. Among the bioactive forms, ABA and SA peaked at 15 DAT in both treatments, but their levels at this time point were higher in the R+Af than in R treatment. IAA and JA levels decreased at 15 DAT, but the decrement of IAA was significantly lower in the R+Af than R treatment. Among CKs, cZ and iP decreased at 15 DAT in both treatments. The levels of DZ and tZ increased moving from 0 to 15 DAT, with DZ levels being higher in R than (in) R+Af treatment at 15 DAT (Fig. 10B).

### *A. filiculoides* and its symbiont released hormonal compounds in the media

*A. filiculoides* affected the contents of the hormonal compounds in the media as significant differences emerged between R and R+Af treatments at 15 DAT (Fig. S9A). Notably, among the compounds that contributed to their separation, there were the phenolics PAAM and SA, along with several auxins, CKs, and GA19 (Fig S9B).

Four of the 35 quantified compounds at 15 DAT were significantly different between the R + Af and R media. Three of these compounds, iP9G, OxIAA and OxIAA-Asp, showed significantly higher levels in the R+Af medium, while the levels of SA significantly decreased (Table S12 and Fig. S10). To test the hypothesis that *T*. *azollae* provided some of the phytohormonal compounds detected in the R + Af medium, phytohormone levels were assessed in the *T. azollae* (Ta) medium after 7 days of growth and compared to the control medium, to which no exogenous hormones were added. The analysis revealed that, in the Ta medium, 4 compounds, namely the CKs iP, cZ and DZOG, and the auxin OxIAA-Glu accumulated (Fig. S11).

## Discussion

### *A. filiculoides* alters the patterns of RSA and the aerial organ development in rice

RSA plasticity is a significant trait enabling plants to cope with abiotic stress (Lavenus *et al*., 2013). Understanding the effectors controlling this trait is therefore crucial to optimizing resource use by crops in a more efficient and sustainable way. The present study shows *A. filiculoides* as a potent trigger of RSA plasticity in rice. A divergent root architecture between co-cultivated and control rice plants becomes in fact macroscopically apparent at 30 DAT, when *A. filiculoides* co-cultivated rice plants show more and shorter ARs than control plants. These changes in AR patterning coincide with a higher number of LRs. The dynamics of root development change with time because at 60 DAT, the ARs of co-cultivated rice become significantly longer and slightly more numerous than those of the control. Thus, *A. filiculoides* shapes the growth and spatial organization of rice roots. Likely, the different RSA exhibited by rice plants in the presence of *A. filiculoides* optimizes nutrient uptake and metabolite exchange between rice and the surrounding aqueous environment.

Co-cultivated rice plants also show increased height and number of tillers from 14 and 21 DAT, respectively. From 28 DAT, the number of leaves is significantly higher in co-cultivated rice. Notably, this number remains higher until the last sampling time point (63 DAT). A higher leaf number in Azolla co-cultivated rice plants was also recently reported by Bazihizina et al. (submitted) after 60 days of hydroponic condition. The same study also reported on a higher fresh shoot biomass and shoot-to-root ratio at this time point in the presence of Azolla. Conversely, under our experimental condition, the rice fresh aerial biomass was significantly higher at 30 DAT when it was also significant the decrease of the root/shoot ratio in R+Af vs R treatment. It is likely that the different experimental set up between present and Bazihizina et al.’s study results in a different timing of rice organ development.

Overall, the present study provides an in-depth analysis on the morphogenic effects that co-cultivated *A. filiculoides* exerts on rice over its early phases of vegetative growth. Because of the experimental set up pursued in the present study, we can also conclude that these effects do neither result from the inorganic nitrogen nor from the VOCs released by the fern. By replacing the growth media frequently and employing only actively growing *A. filiculoides* plants, the content of inorganic nitrogen available to rice co-cultivated plants did not increase compared to the media in which rice plants were grown alone. Yet, *A. filiculoides* co-cultivated and control rice plants were grown side by side in the same growth chamber to allow the VOCs emitted by the fern to diffuse freely among rice plants, regardless of the treatment.

### RSA changes are preceded by a boost of NO and differential expression of iron related genes at 15 DAT

The changes in RSA seen in rice plants co-cultivated with *A. filiculoides* at 30 DAT imply metabolic and molecular changes that occurred at earlier time points. NO, a highly reactive redox signalling molecule, is a central co-regulator in many growth and developmental processes (Qiao and Fan, 2008; Yu *et al*., 2014; Sánchez-Vicente, *et al*., 2019), including the induction and formation of AR and LR (Geiss *et al*., 2009; Xiong *et al*., 2009; Correa-Aragunde *et al*., 2016). Similarly, the reactive nitrogen species (RNS) and ROS signaling molecule network, in synergy with hormonal signaling pathways, control RSA (Prakash *et al*., 2020). The changes in rice RSA observed at 30 DAT are preceded at 15 DAT by a significant increase of NO contents and differential regulation of almost 500 genes in the roots. The GO analysis of DEGs unveiled the commitment of different molecular pathways in response to *A. filiculoides*, and a network hub made up of the S-adenosylmethionine cycle, response to iron, and response to NO (Fig. 7). Thus, transcriptomic data are consistent with the biochemical evidence of higher NO levels at 15 DAT in the roots of rice co-cultivated with *A. filiculoides*.

NR is the most critical source of NO. in plants, and the nitrate concentration/availability in the rooting medium can affect the amount of NO via mediation of NO synthase (NOS) and NO. reductase (NR) activity (Yamasaki *et al*., 1999; Meyer *et al*., 2005; Yamasaki, 2005; Zhao *et al*., 2007) and hence growth. According to Sun and colleagues (2015), NO generated by the NR pathway by increasing LR initiation and the inorganic N uptake rate may represent a strategy for rice plants to adapt to a fluctuating nitrate supply. In keeping with the biochemical quantification of NO. in the roots of co-cultivated rice, genes involved in NO. biosynthesis, *NR2*, and nitrate uptake, such as *NRT1.1B,* were significantly upregulated at 15 DAT, as was the *nitrate and chloride transporter NRT* (Table 2). Additionally, *NR* upregulation was significant from 9 DAT as shown by qRT-PCR analysis. In

Arabidopsis *NRT1.1B* plays multiple roles, one of which is as auxin transporter at low nitrate concentrations (Krouk *et al*., 2010; Forde, 2014). Recently, *NRT1.1B* was found to promote root-to-shoot nitrate translocation and *NRT1.1-NR2* overexpression to improve NUE (Gu and Yang, 2022). The upregulation of *NRT1.1B* in the roots of *A.filiculoides* co-cultivated plants cannot be explained by an increase of nitrate content in the medium. This observation sets the stage to future investigations aimed at identifying additional triggers of *NRT1.1B* regulation. There might be other nitrogen sources present in the media or the different hormonal profiles exhibited by the roots of *A. filiculoides* co-cultivated plants. In this context, we observed that in the presence of the fern, the levels of organic nitrogen in the media, such as small peptides and amino acids, as shown in the companion paper by Consorti *et al*. (2024, Preprint), and those of hormones such as auxin and ABA (see below) increased. Yet, NO. and ABA are interlocking molecules that can exert a combined effect on expression profiles of key genes involved in N-uptake and translocation under stress (Sahay *et al*., 2021).

The regulation of Fe homeostasis is among the key roles of NO (Tewari *et al*., 2021). The effects of NO on Fe homeostasis have been mainly investigated in relation to Fe deficiency, a condition that induces the upregulation of genes for Fe uptake. However, from our transcriptomics data emerged downregulation of such genes. Fe plays a significant role in determining RSA. The availability and uptake of iron have an impact on primary and LR growth and root hair development thereby inducing RSA plasticity (Muller and Schmidt, 2004; Li *et al*., 2016). Also, alternating wet and dry periods in the irrigation of rice leads to the alternant occurrence of Fe deficiency and Fe excess during the rice growth period (Zhang *et al*., 2022). As a consequence, rice has developed a sophisticated mechanism to enhance its Fe stress tolerance and cope with such situations. This mechanism is based on a combination of the chelation-based strategy (Strategy II) and some features of the iron reduction-based strategy (Strategy I) (Li *et al*., 2020). In strategy II, MA family phytosiderophores are synthesized in vesicles and secreted out of the root to chelate Fe^3+^. During the synthesis of DMA, three sequential enzymatic reactions are catalyzed by nicotinamide (NA) synthase (NAS), NA aminotransferase (NAAT), and deoxymugineic acid synthase (DMAS) (Li *et al*., 2020). NAS activity depends on the availability of methionine, which links the S-methionine salvage pathway (MSP) to iron absorption and homeostasis, as it emerged from the ShinyGO analysis (Figures 5, 7).

The MSP pathway is severely downregulated in co-cultivated rice roots. Within this pathway, which is also important for the biosynthesis of isoprenoids and ethylene, the downregulation of *IDI2* emerged from 9 DAT onwards and that of IDI4 from 5 DAT as per qRT-PCR data. After being synthesized in root cells, DMAs are secreted into the rhizosphere by TOM1, whose gene is downregulated in the presence of *A. filiculoides* from 9 DAT (Fig 8). The Fe^3+^-DMA complexes formed in the rhizosphere are transported into root cells by YSL proteins. The rice *YSL2* is significantly downregulated from 9 DAT onwards (Fig.8). Despite preferentially being a strategy II plant, rice also absorbs Fe^2+^ directly via IRTs. In the presence of *A. filiculoides*, both *IRT1* and *2* are severely downregulated at 15 DAT (Table S4). Rice compartmentalizes excessive Fe as ferritins or in vacuoles by inducing the expression of *FER*s and *VITs,* respectively (Zhang *et al*., 2022). Not only *FER1* and *FER2* but also *VIT1-22* and *VITH2* are upregulated in the presence of *A. filiculoides* (Table S4). *FER1* is upregulated from 9 DAT, while *FER2*, the highest expressed ferritin of the two, from 5 DAT onwards (Fig 8). The expression of *FERs* in rice can depend on metals and oxidative stress (Stein *et al*., 2009). In maize *FER2* is regulated through an ABA-dependent pathway whereas *FER1* requires NO and iron through an ABA-independent pathway (Petit *et al*., 2001). We note that both NO and ABA levels are increased in *A. filiculoides* co-cultivated rice roots at 15 DAT. The expression patterns of genes involved in iron absorption, homeostasis, inclusion and sequestration emerged in the present study nicely overlap with those in the roots of rice grown in hydroponics for 14 days under different levels of iron excess (Aung *et al*., 2018). This suggests that *A. filiculoides* co-cultivated rice plants experience a condition of iron excess. However, the increased bioavailability of iron in the presence of *A. filiculoides* seems not to reach toxic levels in rice since no specific morphological symptoms of excess iron, such as bronzing of leaves and roots and stunted roots, occurred.

The downregulation of genes for siderophore biosynthesis in the roots of *A. filiculoides* co-cultivated rice plants might imply that these plants rely on the siderophores released into the medium by the cyanobacteria hosted in the fronds of *A. filiculoides* for iron uptake. Cyanobacteria produce siderophores (Singh, 2014; Chakraborty *et al*., 2019) and those hosted in *Azolla* fronds could make iron and other nutrients more bioavailable in the medium.

Due to the higher iron and nitrate contents in rice roots, a robust generation of ROS (Nguyen *et al*., 2022) and RNS (Tewari *et al*., 2021; Kirk *et al*., 2022) is induced, and various detoxification responses are activated (Aung *et al*., 2018), so plants can quickly adjust growth to the environment by influencing the cellular redox state and signalling (Yuan *et al*., 2013). The significantly higher content of NO also modulates the antioxidant system. In our system, we found the upregulation of several genes involved in the antioxidant system (Table S5) in the roots of *A.filiculoides* co-cultivated plants at 15 DAT.

### *A. filiculoides* induces changes in phytohormone levels in rice roots and growth media

The morphological differences in root patterning along with the evidence that among the DEGs at 15 DAT there are genes related to phytohormone signalling prompted us to analyse the phytohormone levels in rice roots, as well as the hormones present in the cultivation media of rice grown with and without *A. filiculoides*, and those likely supplied by the Azolla symbiont *T. azollae*.

Among the metabolites quantified, only 7 are differentially accumulated between rice roots from R and R+Af treatments at 15 DAT. The levels of two of them, the CK iP7G and the phytoalexin sinAC decreased, while those of the ABA, cZROG, DPA, IAA-Glu and IAA-Asp significantly increased in the presence of *A. filiculoides*. The high levels of ABA in the roots of rice co-cultivated with *A. filiculoides* at 15 DAT is noteworthy. ABA activity is associated, downstream of ethylene action and along with ROS, to ARs initiation in rice (Mhimdi and Pérez-Pérez, 2020). This occurs under waterlogging conditions and directly depends on the continuously produced auxin in the shoot, which is transported through the stem to the roots (Mhimdi and Pérez-Pérez, 2020). Notably, AR and LR initiation and growth are the most noticeable morphological changes exhibited by rice roots in the presence of *A. filiculoides.* In keeping with the increased levels of ABA, there is the upregulation of three ABA responsive genes (*ASR*s) (Table S6). The qRT-PCR analysis showed that *ASR3*, the most highly expressed *ASR*, is upregulated not only at 15 DAT, but also at earlier time points, 9 and 12 DAT (Fig. 8). This finding suggests that the ABA contents increase in rice roots occurred earlier than at 15 DAT. Also, the upregulation of two supposedly negative regulators of the ABA response genes, *PP2C30* and *PP2C27* (Table S6), may indicate a need to control or reduce the effects of ABA to regulate the RSA. In line with this observation, among other differentially expressed *WRKYs*, the marked upregulation of *WRKY40* (FC= 4.20, Table S6) at 15 DAT is noteworthy. In *A. thaliana* this gene is rapidly induced by ABA and acts as a transcriptional repressor of ABA response (Chen *et al*., 2010).

It is generally assumed that CKs act as shoot growth-promoting factors and negative regulators of root development. For instance, exogenous cytokinin treatments inhibit root elongation, but increase plant height and nutrient contents in aerial organs (Beemster and Baskin, 1998; Zahir *et al*., 2001). CKs also inhibit genes related to iron absorption and homeostasis such as *IRT1, FRO2* and *FIT* (Séguéla *et al*., 2008; Gao *et al*., 2019). Thus, along with the supply of siderophores, the provision of CKs by *A. filiculoides* could explain the downregulation of rice genes for iron uptake and the changes in RSA. Yet, the competence of *T. azollae* and its host to synthesize and release CKs in the media coupled with the evidence that genes for CKs biosynthesis are not differentially expressed in rice roots, lead us to argue that rice roots might indeed perceive and even metabolizes exogenous CKs (Zahir *et al*. 2001). Along this reasoning, we note the upregulation of two key genes in CK signaling, the *A-type response regulators RR1* (FC= 2.17) and *RR10* (FC= 2.33) in the roots of *A. filiculoides* co-cultivated plants (Table S6). Genes of G subfamily ATP-binding cassette (*ABCG*) code for CK transporters. In rice only the *OsABCG18* has been characterized as involved in long-distance transport of CKs thus far (Zhao *et al*., 2019). The upregulation of another a *ABCG transporter*, *Os01g0836600* (FC= 2.6,Table S6) in the roots of *A.filiculoides* co-cultivated rice plants paves the way for future research to assess whether this gene could be added to the CKs transporters.

Many aspects of LR formation from priming to emergence are controlled by auxins (Lavenus *et al*., 2013). Although the concentration of IAA increases only slightly in the roots of *A. filiculoides* co-cultivated rice at 15 DAT, that of its storage forms increases significantly. Thus, the presence of the fern impacts on the homeostasis of this hormone. The upregulation at 15 DAT of *NRT1.1,* might concur to shape rice RSA via re-distribution of auxin. This goes along with the marked increase of the auxin responsive gene, *small auxin-up RNA 3* (*SAUR3;* FC = 6.43) and the auxin transporter, *ABCB* (FC = 1.75) (Table S6). Auxin homeostasis is partly sustained by the *GH3* gene family, which can be seen as supervisors of the fluctuation of auxin levels. Although evidence for its function is missing, the increase of the *GH3.12* mRNA levels from 9 DAT onwards may suggest its involvement in the conjugation of amino acids to IAA, so explaining the increase of the IAA storage from IAA-Glu at 15 DAT (Fig 10).

Ethylene is a hormone that plays a vital role in regulating RSA, by acting together with auxin and other phytohormones (Růžička *et al*., 2007; Carvalho *et al*., 2015). We have not quantified the levels of ethylene, however, since two genes involved in ethylene response, *ERF32* (FC = 1.36) and *ERF37* (FC = 3.70), are overexpressed (Table S6), we can infer that the presence of *A. filiculoides* may induce the ethylene signaling pathway in rice roots.

Although no differences in SA levels emerged in the roots of control and *A. filiculoides* co-cultivated plants, their levels decrease in the R+Af media. This observation goes along with the capacity of aquatic organisms such as *Azolla* and *Lemma* spp. to uptake SA and other organic compounds from the media (Maldonado *et al*., 2022) and with the observation that, although some of the key genes for SA biosynthesis seem to be absent in its genome, *A. filiculoides* is responsive to exogenously applied SA (de Vries *et al*., 2018). Being SA a key hormone in plant immunity, lower levels of SA in the medium may indicate that co-cultivation with *A. filiculoides* can perturb the capability of rice or of both plants to interact with each other and with the environment.

### The boost in the development of rice aerial organs reflects the perturbation in the hormonal content in leaves of Azolla co-cultivated rice plants

The morphological changes of aerial organs of *A. filiculoides* co-cultivated plants seem to precede those occurred in roots. Number of leaves, plant height and number of tillers are in fact significantly higher in co-cultivated rice vs control plants well before 30 DAT, when differences in the RSA pattern emerged. However, the differentiation of more organs (tillers and leaves) and, more interestingly, the increase in plant height, seen since 14 DAT, are not sustained at the expenses of root biomass, at least up to 30 DAT. Once again, signal exchanged at root levels between the two partners might have concurred to a different balance of regulators in the first stage of rice aerial organ development. The higher levels of auxins, either bioactive, precursor and storage forms, and of the GA precursor GA19, in the leaves of *A. filiculoides* co-cultivated rice plants suggest this is indeed the case. Auxins in fact promote leaf cell elongation and cell division at leaf node, resulting in an increase in the leaf pitch and gibberellins have a central role in determining leaf growth and height (Zhao *et al*., 2021; Sprangers *et al*., 2020). While the levels of bioactive CK forms do not increase in the leaves of *A. filiculoides* co-cultivated vs to control rice plants, those of the CK precursor, transport and storage forms are higher. Thus, in the presence of *A. filiculoides* the homeostasis of CKs in rice leaves is perturbed. Because CKs regulate the expression of nitrogen transporters from old to new leaves (Kiba *et al*., 2011), it would be interesting to assess whether the gradient of nitrogen distribution in rice canopy changes following co-cultivation with *A. filiculoides*.

In the leaves of *A. filiculoides* co-cultivated plants the levels of ABA are also higher. ABA is known to affect mainly seed dormancy and germination, stomatal closure and stress tolerance. However, the evidence that ABA-deficient mutants in *A. thaliana* show smaller plant stature and leaf growth than wild type suggests that endogenous physiological concentrations of ABA may act as growth promoter. In line with this, these mutants exhibit reduced cell area and cell number (Horiguchi *et al*., 2006; Barrero *et al*., 2005), while in wild type *A. thaliana* plants ABA maintains shoot development and leaf expansion in well-watered plants (LeNoble *et al*., 2004). It is therefore conceivable that as in *A. thaliana* ABA also has a dual function in rice: growth inhibitor under stress and growth promoter under controlled conditions (Cheng *et al*., 2002).

Besides its function during biotic and abiotic stress, SA plays a crucial role in the regulation of physiological and biochemical processes during the entire lifespan of the plant (Rivas-San Vicente and Javier Plasencia, 2011). Although a combination of several internal and external stimuli contributes to plant growth and development, it was shown that micromolar application of SA to seedling/ plantlet shoots of different species increases stem diameter, leaf number and fresh biomass (Tucuch-Haas *et al*., 2017). Thus, by affecting leaf and chloroplast structure, SA is an important regulator of photosynthesis. As an example, the photosynthetic rate is increased in maize sprayed with 10−2M SA (Khodary, 2004). In turn, it is likely that the higher levels of SA detected in the leaves of Azolla co-cultivated plants might sustain a higher photosynthetic rate and, in turn, the higher areal biomass observed at 30 DAT. Moreover, the higher levels of SA in *A. filiculoides* co-cultivated plants leaves let us argue that treated plants could cope better with stress than control plants.

## Conclusions

Here we show for the first time the morphogenic effects that *A. filiculoides* exerts on rice plants at the early phases of co-cultivation. The alteration in the development program triggered by the fern on rice plants reflects and likely results from the increased availability in the co-cultivation medium of small peptides, lipids and flavonoids, as shown in the companion paper by Consorti *et al*. (2024, Preprint), as well as from the change in hormonal balance in rice roots and leaves, and an increased availability of mineral nutrients. The present study shows in fact that *A. filiculoides* and its symbiont *T. azollae* are competent to release in the liquid medium CKs and storage forms of auxins that are likely transduced and metabolized by rice roots. Moreover, although the presence of *A. filiculoides* induces the upregulation of rice root genes for iron sequestration and compartmentalization, as it occurs when plants experience iron toxicity, co-cultivated rice plants do not show symptoms of iron excess. No lastly, the co-cultivation with *A. filiculoides* induces an increase in SA levels in rice leaves. This observation, along with enhanced formation of ARs and the boost in the development of the above-ground organs in rice seedlings, point towards the co-cultivation with *A. filiculoides* as a sustainable strategy to help rice plantlets coping with abiotic and biotic stress. Thus, future investigations will be carried out for assessing how and to what extent Azolla co-cultivation impacts rice plant development and seed yield under stressful conditions. Finally, the present study highlights a metabolic hub in which different hormones and increased levels of iron and NO likely play a major role in determining a more efficient rice RSA. The employment of rice mutants, impaired in the biosynthesis and perception of these compounds, and different rice varieties, will allow us to understand more about the intricate below-ground metabolic crosstalk taking place between Azolla and rice.

## Abbreviations

SDGs: Sustainable Development Goals
N: nitrogen
NUE: nitrogen use efficiency
PAR: photosynthetically active radiation
N2: atmospheric nitrogen
VOCs: volatile organic compounds
RSA: root system architecture
½MS: half-strength MS nutrients
R treatment: boxes containing sole rice plants (control)
R+Af treatment: rice co-cultivated with *A. filiculoides*
DAT: Days After Transplanting
ARs: adventitious roots
LRs: lateral roots
NH_4_^+^: ammonium
NO_2_^-^: nitrite ion
NO: nitric oxide
DAF-2DA: 4,5-diaminofluorescein diacetate
CTCF: corrected total cell fluorescence
NED: N-(1-naphthyl) ethylenediamine dihydrochloride
DEGs: differentially expressed genes
SPE: solid-phase extraction
IS: internal standards
ROS: reactive oxygen species
RNS: reactive nitrogen species.

## Acknowledgments

This work is dedicated to the memory of our eminent colleague Stefania Pasqualini who conceived this study, earned a grant that supported it, and supervised the doctoral research activity of S.C. We thank Agnese Bertoldi for her hep with plant growth and sampling.

## Author contributions

FP and MK designed the study; SC and AC carried out the entire experimental set. SC, AC, MC and LR performed morphological analysis. SC, CP, MCV, AS, VG, GCZ, PID, MMK and FP performed molecular analyses. LR, PID, MMK and FP supervised the study. SC, LR and FP wrote the manuscript with the contribution of all the authors.

## Data availability

The RNAseq data are available in the NCBI Gene Expression Omnibus database (https://www.ncbi.nlm.nih.gov/geo), accession number GSE278294. All other relevant data can be found within the manuscript and its supplementary data online.

## Supplementary data

**Table S1 -** DEGs and housekeeping genes investigated by qRT-PCR analysis.

**Table S2 -** Levels of inorganic nitrogen forms in growth media at 10, 20 and 30 DAT.

**Table S3 -** DEGs of the methionine savage pahway (MPS).

**Table S4 -** DEGs involved in iron absorption and homeostasis.

**Table S5 -** DEGs related to ROS scavenging/homeostasis.

**Table S6 -** DEGs involved in hormone processing and signalling.

**Table S7 -** Compounds investigated by targeted-hormonomics.

**Table S8 -** Detected and quantified hormonal compounds in the roots of rice grown in the R+Af compared to the R treatment at 15 DAT.

**Table S9 -** Detected and quantified hormonal compounds in the roots of rice grown in the R and R+ Af treatments over time, 0 and 15 DAT.

**Table S10 -** Detected and quantified hormonal compounds in the leaves of rice grown in the R+Af compared to the R treatment at 15 DAT.

**Table S11 -** Detected and quantified hormonal compounds in the leaves of rice grown in the R and R+Af treatments over time, 0 and 15 DAT.

**Table S12 -** Detected and quantified hormonal compounds in the media of rice grown in the R+Af compared to the R treatment at 15 DAT.

**Figure S1 -** Effect of *A. filiculoides* on the biomass of the root apparatus of rice at 15, 30 and 60 DAT. Bar Graphs represent mean values ± SE of control rice (R) and rice grown with *A. filiculoides* (R + Af). P > 0.05.

**Figure S2 -** Effect of *A. filiculoides* on the relative amount of NO in the ARs of rice at 15 and 30 DAT quantified via Griess assay. Bar Graphs represent mean values ± SE of control rice (R) and rice grown with *A. filiculoides* (R+Af) treatments. *, p < 0.05.

**Figure S3 -** Effect of *A. filiculoides* on the above ground part and total biomass of rice (also refer to Fig. 12) at 15, 30 and 60 DAT. Bar Graphs represent mean values ± SE of control rice (R) and rice grown with *A. filiculoides* (R+Af). *, P < 0.05; **, P < 0.01.

**Figure S3 -** Plots of the fold enrichment of the upregulated DEGs. A: BP and B: MF. BP: biological process; MF: molecular function.

**Figure S5 -** Plots of the fold enrichment of the downregulated DEGs. A: BP and B: MF. BP: biological process; MF: molecular function.

**Figure S6 -** sPLSDA (A) and sPLSDA corresponding loadings plot (B) of the most discriminant phytohormones explaining replicates’ distribution in the roots of rice grown with (R+Af) and without *A. filiculoides* (R) at 15 DAT. The higher the Loadings value on the x-axis, the more discriminant the compound (B).

**Figure S7 -** sPLSDA (A) and sPLSDA corresponding loadings plot (B) of the most discriminant compounds explaining replicates classification in the leaves of rice grown in the R+Af and R treatments at 15 DAT.

**Figure S8 -** Significantly different levels of hormonal compounds in the leaves of rice grown with and without *A. filiculoides* at 15 DAT. Unpaired two samples t-test for all nineteen compounds; P < 0.01; FDR cut-off ≤ 0.001. See also Table S12.

**Figure S9 -** sPLSDA (A) and corresponding loadings plot (B) of the most discriminant phytohormones explaining treatments’ variance in the media in the R+Af and R treatments at 15 DAT.

**Figure S10 –** Significant different levels of hormonal compounds in the R+Af and R media at 15 DAT. Unpaired two samples t-test for all four compounds; P < 0.05. See also Table S11.

**Figure S11 -** Differential levels of hormonal compounds in the control and *T. azollae*-growing (Ta) media after 7 days of growth.

