## Supplementary figures S1-S11 for "Co-cultivating rice plants with *Azolla filiculoides* modifies root architecture and timing of developmental stages"

**Fig. S1:** Effect of *A. filiculoides* on the biomass of the root apparatus of rice at 15, 30 and 60 DAT. Bar Graphs represent mean values ± SE of control rice (R) and rice grown with *A. filiculoides* (R+Af). P > 0.05.

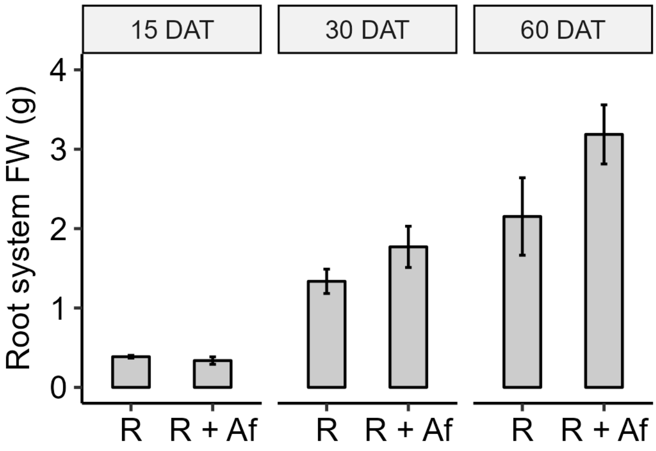


**Fig. S2:** Effect of *A. filiculoides* on the relative amount of NO in the ARs of rice at 15 and 30 DAT quantified via Griess assay. Bar Graphs represent mean values ± SE of control rice (R) and rice grown with *A. filiculoides* (R+Af) treatments. *, p < 0.05.

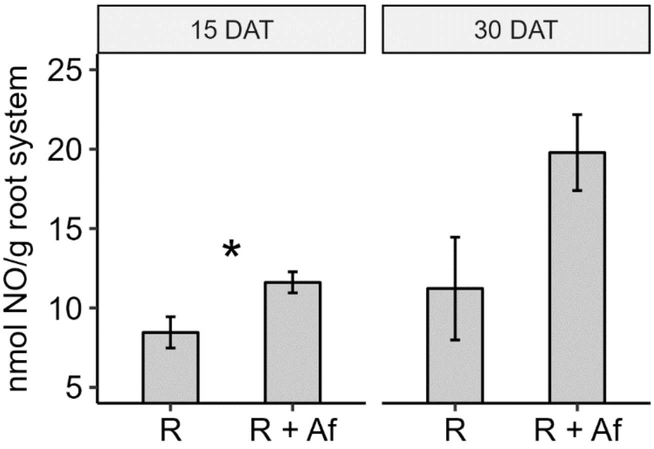


**Fig. S3.** Effect of *A. filiculoides* on the above ground part and total biomass of rice (also refer to Fig. 12) at 15, 30 and 60 DAT. Bar Graphs represent mean values ± SE of control rice (R) and rice grown with *A. filiculoides* (R+Af). *, P < 0.05; **, P < 0.01.

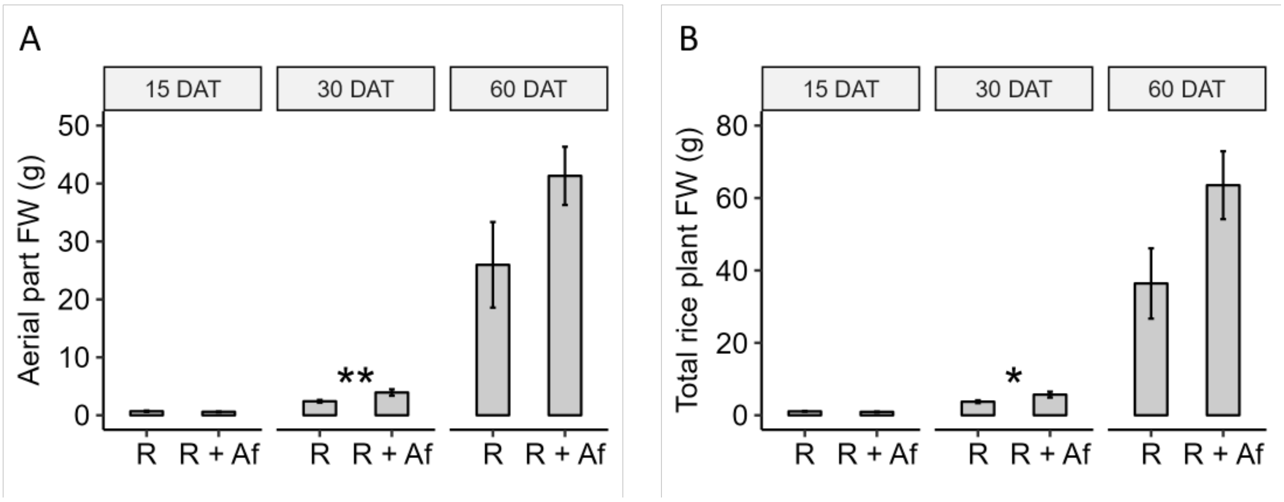


**Fig. S4**. Plots of the fold enrichment of the upregulated DEGs. A: BP and B: MF. BP: biological process; MF: molecular function.

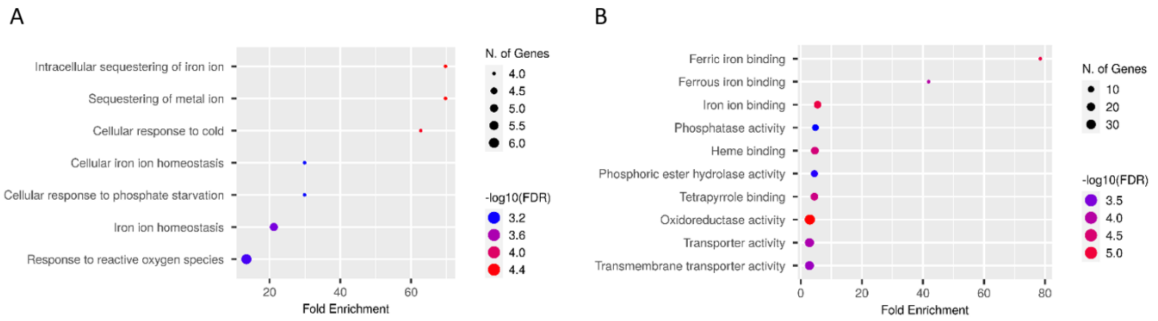


**Fig. S5**. Plots of the fold enrichment of the downregulated DEGs. A: BP and B: MF. BP: biological process; MF: molecular function.

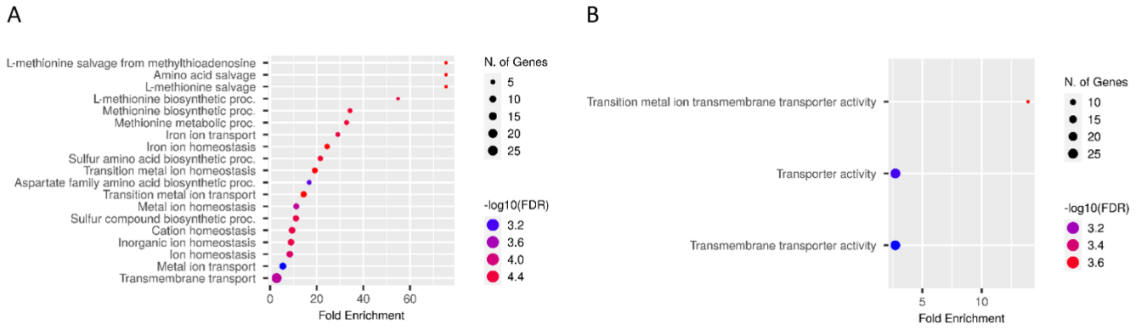

**Fig. S6** sPLSDA (A) and sPLSDA corresponding loadings plot (B) of the most discriminant phytohormones explaining replicates' distribution in the roots of rice grown with (R+Af) and without *A. filiculoides* (R) at 15 DAT. The higher the Loadings value on the x-axis, the more discriminant the compound (B).

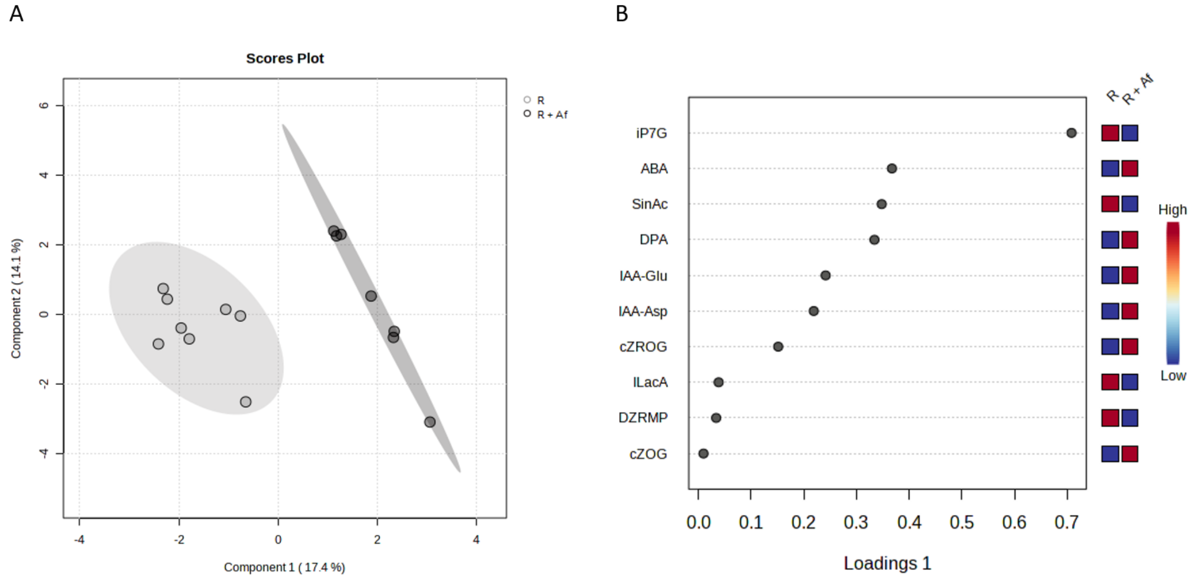


**Fig. S7**. sPLSDA (A) and sPLSDA corresponding loadings plot (B) of the most discriminant compounds explaining replicates classification in the leaves of rice grown in the R+Af and R treatments at 15 DAT.

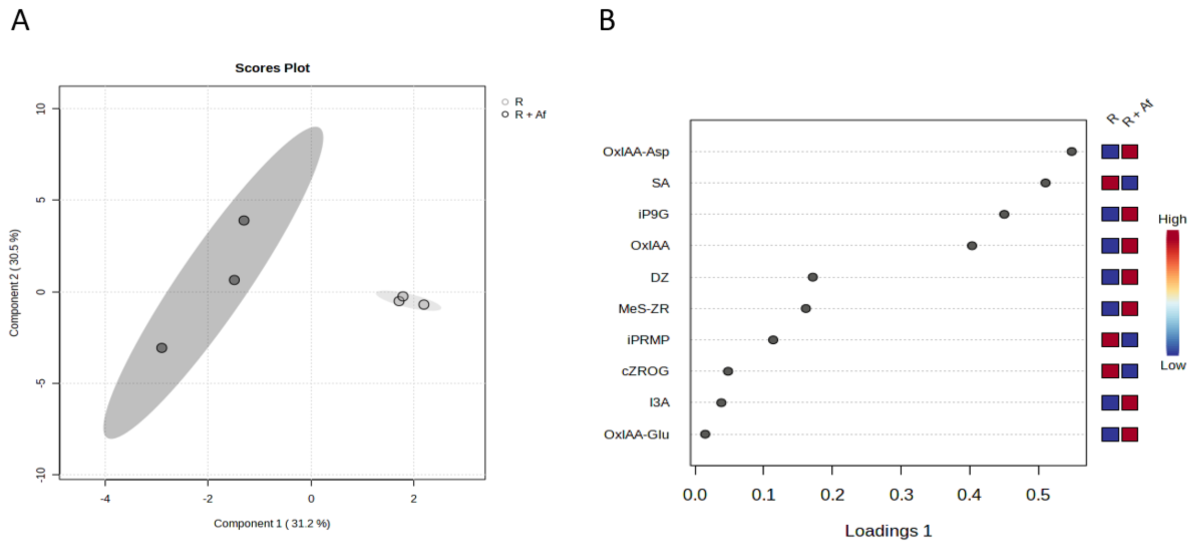

**Fig. S8**. Significantly different levels of hormonal compounds in the leaves of rice grown with and without *A. filiculoides* at 15 DAT. Unpaired two samples t-test for all nineteen compounds; P < 0.01; FDR cut-off ≤ 0.001. See also Table S12.

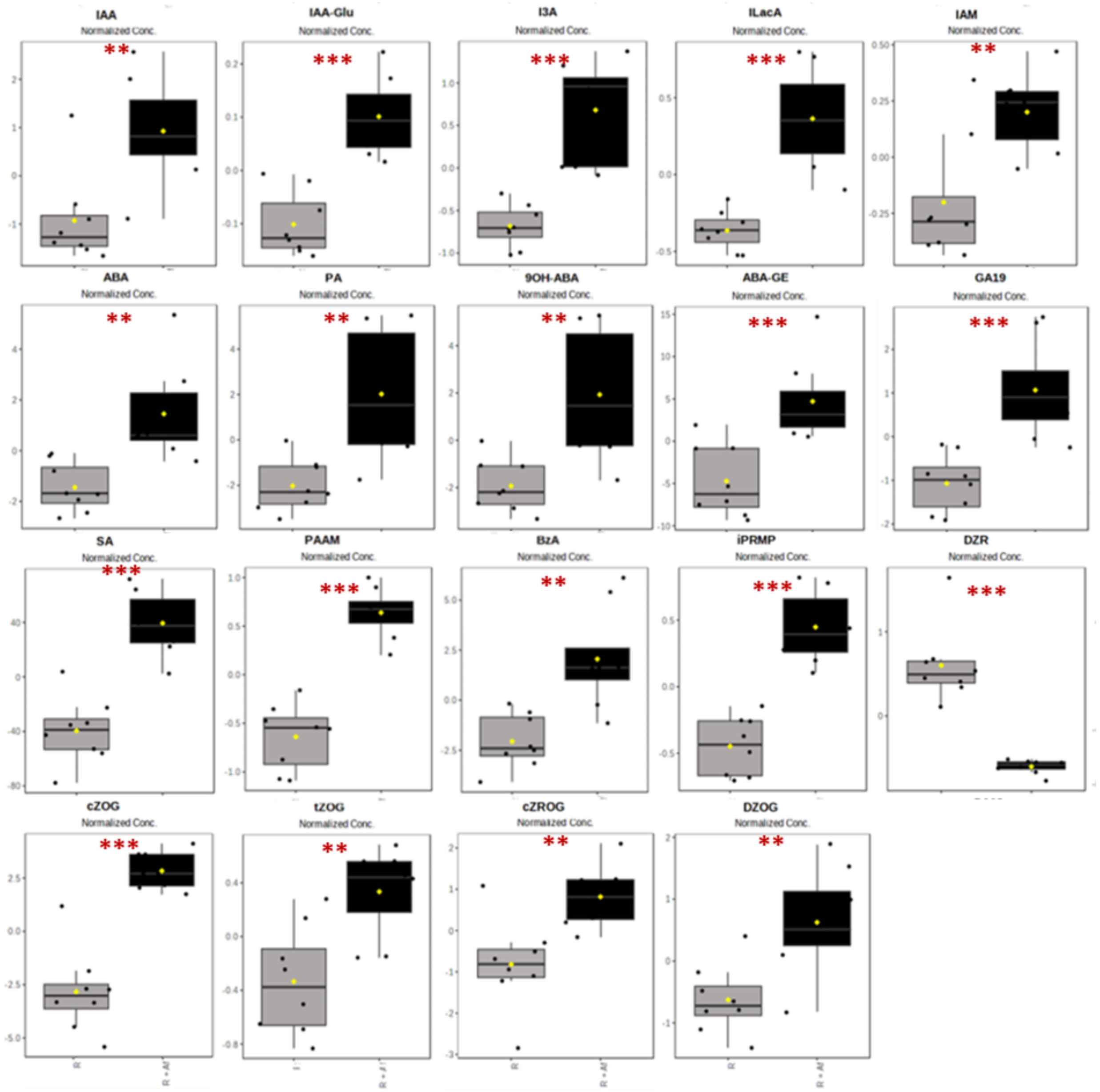


**Fig. S9** sPLSDA (A) and corresponding loadings plot (B) of the most discriminant phytohormones explaining treatments' variance in the media in the R+Af and R treatments at 15 DAT. The higher the Loadings value on the x-axis, the more discriminant the compound.

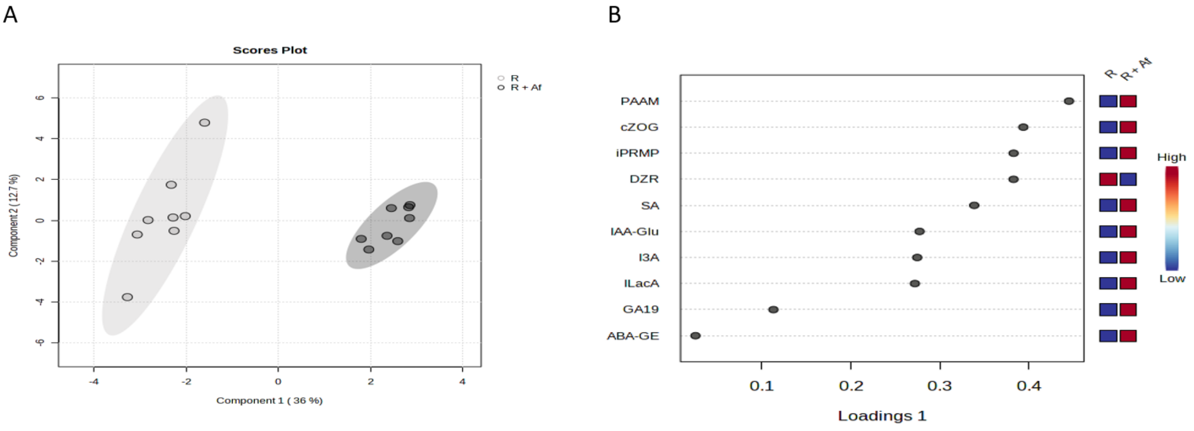


**Fig. S10**. Differential levels of hormonal compounds in the control and *T. azollae*-growing (Ta) media after 7 days of growth.

**Fig. S10**. Significant different levels of hormonal compounds in the R+Af and R media at 15 DAT. Unpaired two samples t-test for all four compounds; $P < 0.05$. See also Table S11.

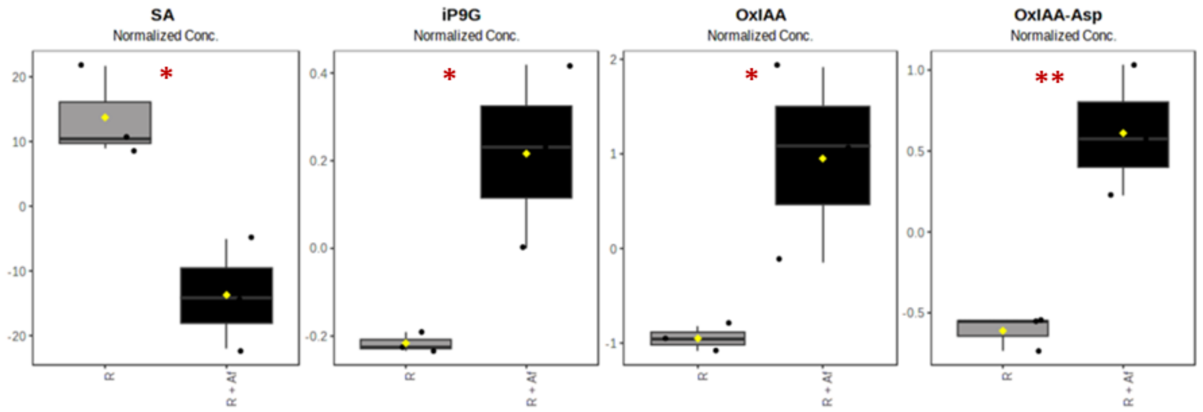


**Fig. S11**. Differential levels of hormonal compounds in control and *T. azollae*-growing (Ta) media after 7 days of growth.

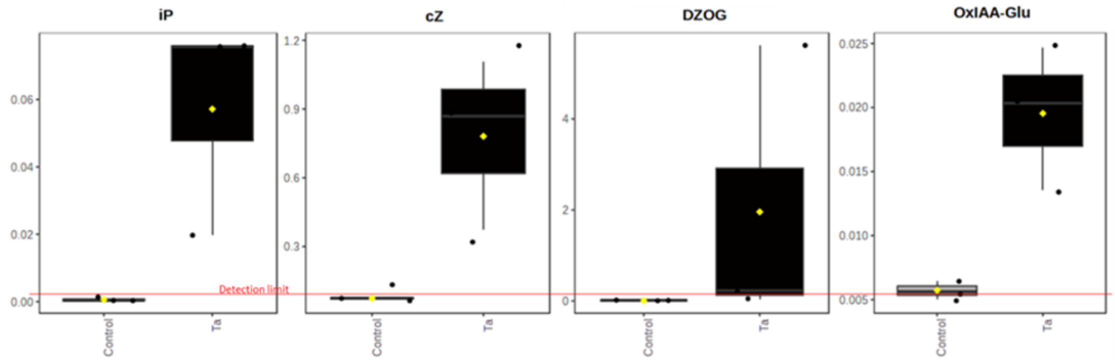
